# TSA-Seq reveals a largely “hardwired” genome organization relative to nuclear speckles with small position changes tightly correlated with gene expression changes

**DOI:** 10.1101/824433

**Authors:** Liguo Zhang, Yang Zhang, Yu Chen, Omid Gholamalamdari, Yuchuan Wang, Jian Ma, Andrew S. Belmont

## Abstract

Genome-wide mapping of chromosomal distances relative to nuclear compartments using TSA-Seq suggests a more deterministic relationship between intranuclear gene position and expression as a function of nuclear speckle distance than radial position. Gene activity increases overall with decreasing distance to nuclear speckles, with active chromosomal regions forming the apex of chromosome loops protruding from the nuclear periphery into the interior. Interestingly, genomic distances to the nearest lamina-associated domain are larger for loop apexes mapping very close to nuclear speckles, suggesting the possibility of genomic “hardwiring” and conservation of speckle-associated regions. To facilitate comparison of genome organization relative to nuclear speckles in human K562, HCT116, HFFc6, and H1 cell lines, here we describe reducing the required cell number 10-20-fold for TSA-Seq by deliberately saturating protein-labeling while preserving distance mapping by the still unsaturated DNA-labeling. Surprisingly, in pair-wise cell line comparisons, only ∼10% of the genome shows a statistically significant shift in relative nuclear speckle distances. These modest shifts in nuclear speckle distance, however, tightly correlate with changes in cell-type specific gene expression. Similarly, half of all loci that contain induced heat-shock protein genes appear pre-positioned close to nuclear speckles, with the remaining showing small shifts towards speckles with transcriptional induction. Speckle association together with chromatin decondensation correlates with expression amplification upon *HSPH1* activation. Our results demonstrate a largely “hardwired” genome organization and specific genes moving small mean distances relative to speckles during cell differentiation or physiological transition, suggesting an important role of nuclear speckles in gene expression regulation.

## Introduction

The mammalian genome is organized non-randomly in the nucleus, with the observed spatial positioning of genes highly correlated with their timing of DNA replication and expression (Ferrai et al. 2010; Meldi and Brickner 2011; Bickmore and van Steensel 2013). A radial distribution of genome organization with a gradient of gene expression increasing from nuclear periphery to interior has been demonstrated in multiple tissue types across a wide range of metazoans (Takizawa et al. 2008b; Bickmore 2013). However, this radial positioning and its correlation with gene expression is highly stochastic. Within a cell population, a given allele can vary widely in its radial position; similarly, different genes with similar transcriptional activity can vary widely in their mean radial positions (Takizawa et al. 2008b). Thus the functional significance of radial gene positioning remains unclear (Takizawa et al. 2008b).

Subsequent measurement of DNA ligation interaction frequencies using Hi-C suggested that the genome could be divided into two major compartments, “B” and “A” (Lieberman-Aiden et al. 2009), which corresponded closely to Lamina-Associated Domains (LADs) and inter-LADs (iLADs) (Kind et al. 2015; van Steensel and Belmont 2017), as well as late-replicating versus early-replicating chromosome regions, respectively (Ryba et al. 2010). These discoveries instead suggested a binary division of the genome into peripherally localized heterochromatin and interiorly localized euchromatin. More recently, higher resolution Hi-C further subdivided the genome into two A subcompartments and three primary B subcompartments (Rao et al. 2014), while newer genomic methods such as SPRITE (Quinodoz et al. 2018) and GAM (Beagrie et al. 2017) both suggested multi-way interactions occurring more frequently in the nuclear interior, specifically at nucleoli and nuclear speckles (Quinodoz et al. 2018). Thus, in addition to the nuclear lamina, these newer results thus point to additional nuclear bodies/compartments shaping nuclear genome architecture, consistent with significant previous literature derived from microscopy analysis of the positioning of specific chromosome loci relative to nuclear speckles, nucleoli, and chromocenters, among other nuclear bodies (Shopland et al. 2003; Wang et al. 2004; Pederson 2011; Politz et al. 2013; van Steensel and Belmont 2017; Jagannathan et al. 2018).

In particular, nuclear speckles, previously defined as interchromatin granule clusters by electron microscopy (Fakan and Puvion 1980), have alternatively been proposed to act as storage sites for factors primarily related to RNA processing or as gene expression hubs for a significant subset of active genes, as reviewed recently (Chen and Belmont 2019). These two functions are not necessarily mutually exclusive. Indeed, an intricate coupling between transcription, RNA processing, and RNA export is now appreciated (Galganski et al. 2017). Nuclear speckles are known to be enriched in factors related to transcriptional pause release, RNA capping and splicing, poly-adenylation and cleavage factors, and RNA export (Saitoh et al. 2004; Galganski et al. 2017). Heuristically, it is easy to imagine how localization of genes near storage sites for all these factors might reasonably lead to increased gene expression. Indeed, using model transgene systems, live-cell imaging has demonstrated an active, nuclear-actin dependent mechanism for moving genes to nuclear speckles over distances up to several microns (Khanna et al. 2014). More recently, both speckle-associated *HSP70* transgenes and endogenous gene loci after heat-shock showed ∼3-50-fold higher nascent transcript levels versus alleles not in contact with nuclear speckles, while live-cell imaging revealed increases (decreases) in nascent transcripts within several minutes after nuclear speckle association (separation) (Kim et al. 2020).

Evidence for nuclear speckles serving as a gene expression hub, however, has been based largely on low-throughput microscopy of a small number of gene loci (Xing et al. 1995; Shopland et al. 2003; Moen et al. 2004; Hall et al. 2006; Hu et al. 2009; Hu et al. 2010). To determine the true prevalence of gene association with nuclear speckles, we developed a novel method to map spatial chromosomal distances relative to a specific nuclear compartment/body (Chen et al. 2018). Based on Tyramide Signal Ampification (TSA) (Bobrow et al. 1989; Raap et al. 1995), which amplifies immunostaining through generation of tyramide free radicals by horse-radish peroxidase (HRP), TSA-Seq exploits the exponential decay gradient of tyramide labeling from a point source to convert sequencing reads into a “cytological ruler”, reporting on mean chromosomal distances from immuno-stained nuclear compartments (Chen et al. 2018). Applying TSA-Seq to K562 cells revealed a nuclear lamina to speckle axis correlating with a gradient in gene expression levels and DNA replication timing, among other genomic features. Further analysis revealed this overall gradient in gene expression levels with distance to nuclear speckles represented the superposition of Hi-C A1 subcompartment active regions localized very close to nuclear speckles and A2 subcompartment active regions localized at intermediate speckle-distances. More specifically, active chromosomal regions formed the apex of chromatin loops protruding into the nuclear interior from the nuclear periphery, with Hi-C A1 active regions localizing significantly closer to speckles than Hi-C A2 active regions (Chen et al. 2018). Interestingly, distances in DNA base-pairs to the nearest lamina-associated domain were larger for loop apexes mapping close to nuclear speckles, suggesting the possibility of genomic “hardwiring” of speckle-associated regions and predicting speckle-associated regions might be conserved in different cell types.

To test this “hardwiring” model of genome organization relative to nuclear speckles, here we compare relative nuclear speckle distances genome-wide in four human cell lines and changes within one cell line after heat shock. A major limitation of the original TSA-Seq protocol was its limited sensitivity, requiring ∼100-1000 million cells per replicate. To facilitate these comparisons, we therefore first increased the sensitivity of TSA-Seq 10-20-fold through a deliberate “super-saturation” staining approach.

Genome organization relative to speckles was highly conserved, with ∼90% of the nuclear genome maintaining similar relative distances across quite different cell types ranging from human ESCs to fibroblasts. This contrasts sharply with the ∼50% of changes in the LAD versus inter-LAD organization encountered in these same cell type comparisons of DamID mapping (unpublished data from van Steensel group (NKI): https://data.4dnucleome.org) and ∼30% of changes in mouse cell type comparisons (Meuleman et al. 2013). However, a small fraction of the genome, divided into hundreds of domains several hundred Kbps to several Mbps in size, show small but statistically significant shifts relative to nuclear speckles ranging from an estimated 0.1-0.4 microns in mean distance. These shifts highly correlate with significant changes in gene expression levels and histone marks, with regions moving closer to nuclear speckles enriched in genes with cell-type specific functions. Moreover, examination of the heat-shock response revealed “hardwiring” of ∼half of all heat-shock induced loci to be already in close contact with nuclear speckles, with the remaining heat-shock induced loci positioned at intermediate distances to nuclear speckles but all moving closer after heat-shock induction of gene expression. Combined DNA and RNA immune-FISH of the HSPH1 locus revealed increased speckle association, accompanied by chromatin decondensation, was correlated with an ∼2-3-fold gene expression amplification of HSPH1 nascent transcript levels.

Our results suggest that cells tightly regulate their genome’s organization relative to nuclear speckles and a close relationship between changing gene expression and distance to nuclear speckles for a substantial fraction of genes.

## Results

### Greater than 10-fold increased sensitivity using “super-saturation” TSA-Seq

A major experimental limitation of our original TSA-Seq protocol was its requirement for large cell numbers (Chen et al. 2018). Although Tyramide Signal Amplification (TSA) is typically described as labeling the tyrosine residues of proteins, core histones showed very low TSA-labeling (by western blotting), presumably due to inaccessibility of their tyrosine residues. This poor histone labeling suggested that TSA chromatin labeling would be heavily biased towards DNA regions with high, tyrosine-rich nonhistone protein content. In contrast, we discovered that tryamide free-radicals label DNA uniformly across the genome (Chen et al. 2018). TSA-Seq, therefore, was based on tyramide-labeling of DNA, even though the low efficiency of TSA DNA labeling translates into a requirement for large cell numbers-anywhere from ∼100-1000 million cells per TSA-replicate, depending on the staining target and desired spatial resolution. This requirement for such large cell number prevents the routine use of the original TSA-Seq method to investigate genome organization.

During our development of TSA-Seq (Chen et al. 2018), we explored a wide range of tyramide concentrations and reaction times, choosing staining conditions which maximized signal strength while maintaining a high signal to nuclear background ratio. An obvious solution to increasing the DNA labeling after TSA would be to increase the tyramide concentration and/or TSA reaction time, given that tyramide free-radicals are produced by an enzymatic reaction. Curiously, though, TSA staining of nuclear speckles with increasing tyramide concentrations and reaction times appeared to produce increasing nonspecific staining, spreading throughout the nucleus and even cytoplasm. This unexplained increase in nonspecific staining after increasing production of tryamide free-radicals appeared to prevent increasing TSA-Seq sensitivity by increasing tyramide concentrations and/or TSA reaction times.

Upon further consideration, however, we realized that the TSA labeling visualized by light microscopy after anti-biotin staining was likely dominated by the protein, rather than nucleic acid, biotin-tyramide labeling, given the observed low level of DNA TSA labeling revealed by molecular analysis. We hypothesized that progressive saturation of protein tyrosines causes non-specific TSA labeling to spread progressively outwards from nuclear speckles, with the weaker DNA tyramide labeling, far from saturation and still specific for the staining target, masked by this stronger protein labeling.

To test these hypotheses, we first designed a series of TSA labeling conditions with increased tyramide-biotin concentrations and reaction times (Conditions A-E) using anti-SON nuclear speckle TSA labeling in K562 cells (Fig. 1A, Fig. S1A). Condition A corresponds to our original, “TSA-Seq 1.0” conditions (Chen et al. 2018). Nuclear speckle TSA labeling visualized by light microscopy increased from Conditions A to C but plateaued, with increases in background staining, from Conditions C to E (Fig. 1A-B). Condition E produced higher tyramide-biotin labeling in the cytoplasm with no visible specific nuclear speckle staining above the nucleoplasmic staining. To then reveal the hypothesized, nonsaturated and specific DNA TSA-labeling above the background of saturated protein TSA-labeling, we “blocked” protein tyrosine groups using high concentrations of tyramide-biotin (Condition E), and then performed a second TSA using tyramide-FITC (Condition A) (Fig. 1C). As expected, TSA labeling using the under-saturating Condition A produced similar nuclear speckle staining after both rounds of TSA labeling (Fig. 1D, top). As predicted by our hypothesis (Fig. 1C), while Condition E TSA labeling produced diffuse biotin-labeling throughout the cell, a 2^nd^ TSA labeling (Condition A) revealed nuclear speckle-specific labeling in K562 (Fig. 1D, bottom) and HFFc6 cells (Fig. S1B, bottom).

**Figure 1.**
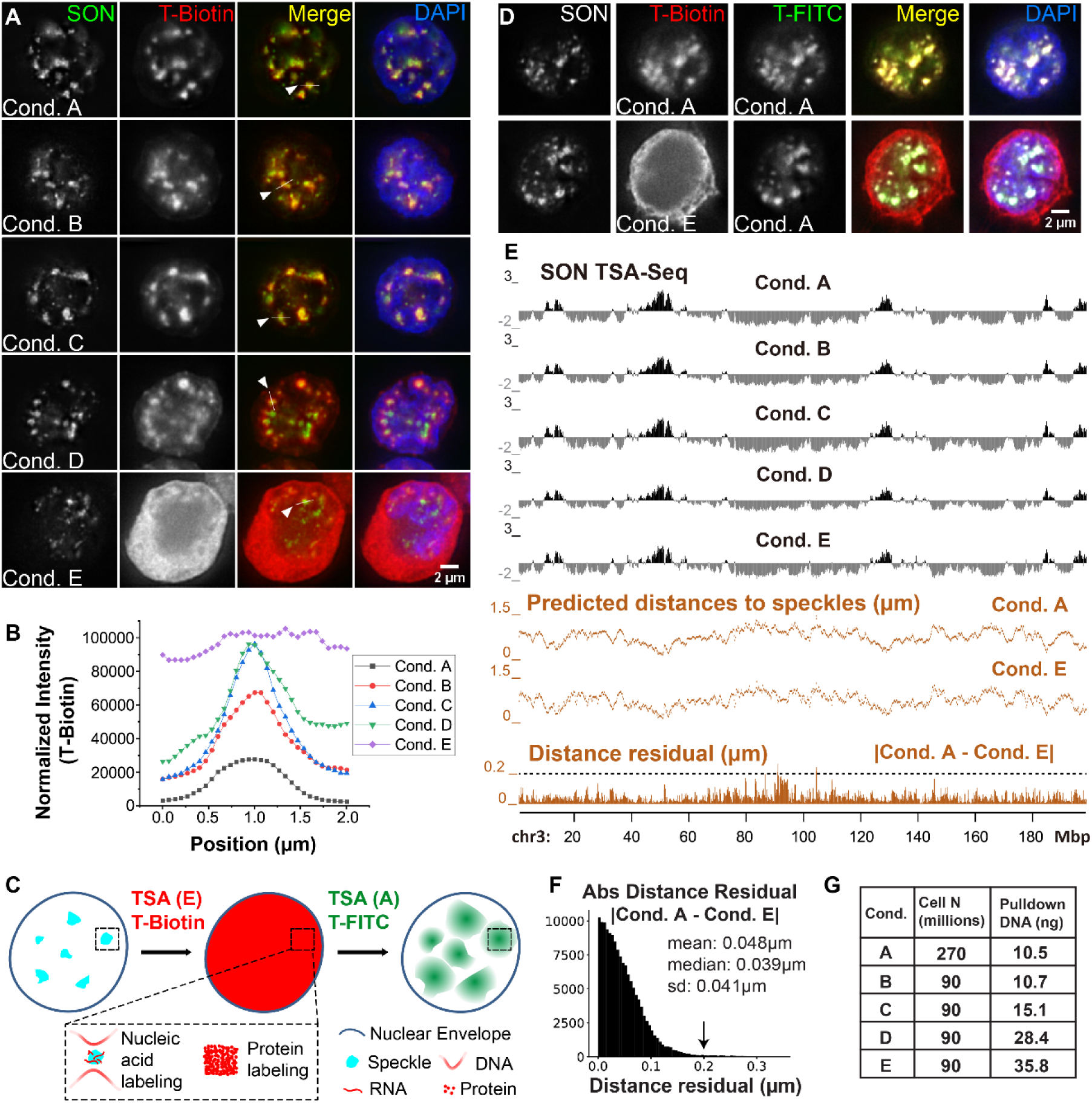
TSA-Seq 2.0 enables 10-20-fold increase in sensitivity but preserves distance mapping capability. **A)** TSA Conditions A-E (K562 cells) show varying nuclear speckle specificity. SON immunostaining of speckles (green), strepavidin tyramide-biotin staining (red), merged channels, plus DNA (DAPI, blue). **B)** Tyramide-biotin intensities along line profiles spanning nuclear speckles in **A)** for Conditions A-E. **C)** Schema predicting results from following Condition E tyramide-biotin TSA staining with Condition A tyramide-FITC TSA staining, assuming Condition E saturates protein but not DNA tyramide-labeling. **D)** Experimental results for schema in **C)**. Top row: control showing two consecutive rounds of Condition A (non-saturating) TSA-labeling using tyramide-biotin and then tyramide-FITC. Bottom row: Same as top row but using Condition E for 1st TSA-labeling. SON immunostaining (grey), tyramide-biotin (red), tyramide-FITC (green), merged channels, plus DAPI (blue). **E)** SON TSA-Seq mapping results for Conditions A-E showing TSA-Seq enrichment scores (black tracks), estimated speckle distances (Conditions A and E, middle orange tracks), and residuals (absolute magnitude) between Conditions A and E distances (bottom, orange track). **F)** Histogram of distance residuals in **E)**: number (y-axis), residual value (x-axis). **G)** Cell numbers used and pulldown DNA yields for TSA Conditions A-E.

We estimated average tyramide-biotin labeling increased from ∼1 biotin /200 kb using Condition A to ∼ 1 biotin/7.5 kb using Condition E (Fig. S1C), predicting ∼1 biotin/0.94 kb over the ∼8 fold-enriched speckle TSA-Seq peaks. Sonicating DNA to ∼100-600 bps ensures few DNA fragments with multiple biotins, maintaining linearity between TSA labeling and pulldown read number.

Speckle TSA-Seq maps generated from Conditions A-E were both qualitatively and quantitatively similar (Fig. 1E). Converting TSA-Seq signals to predicted nuclear speckle distances using exponential fitting of immuno-FISH data (Supplementary Table 1 and 2, Methods) demonstrated similar distances estimated from TSA-Seq data across Conditions A-E (Fig. 1E, Fig. S2A). Distance residuals across the genome between Conditions A and E were nearly all smaller (<0.05 µm mean and median) than the microscopy diffraction limit of ∼0.25 µm (Fig. 1E-F, Fig. S2). Conditions A, C, and E also produced similar SON TSA-Seq maps in HCT116 cells (Fig. S3). Thus using Condition E we were able to increase pulldown DNA yields up by 10-20 fold, relative to TSA-Seq 1.0 (Condition A) (Fig. 1G, Fig. S3D), reducing required cell numbers to ∼10-15 million (to obtain ∼5 ng pulldown DNA), without significant quantitative changes in nuclear speckle distance estimations.

### Speckle-associated domains remain largely conserved, but small, statistically significant changes highly correlate with gene expression changes

Previously, we defined chromosome regions with top 5% nuclear speckle TSA-Seq scores as SPeckle-Associated Domains (SPADs) as they were near-deterministically positioned near nuclear speckles, with mean estimated speckle distances of <0.32 µm and ∼100% of alleles positioned within 0.5 µm of speckles by FISH (Chen et al. 2018). To test whether SPADs are conserved among different cell types, we applied SON TSA-Seq to four cell lines (Supplementary Table 3, Fig. 1A, Fig. S3A, S4, Fig. 2A), each with two biological replicates: K562 erythroleukemia cells, HCT116 colon carcinoma cells, HFFc6 human foreskin fibroblast cells immortalized by hTERT expression, and H1 human embryonal stem cells (hESCs). Our new supersaturation TSA labeling conditions (“TSA-Seq 2.0”), greatly facilitated extension of SON TSA-Seq to these cell lines. Multiple batches of TSA 1.0 staining, each done with many flasks of cells and repeated over multiple weeks, were replaced with a single TSA-Seq 2.0 staining, performed with 1-3 flasks of cells.

**Figure 2.**
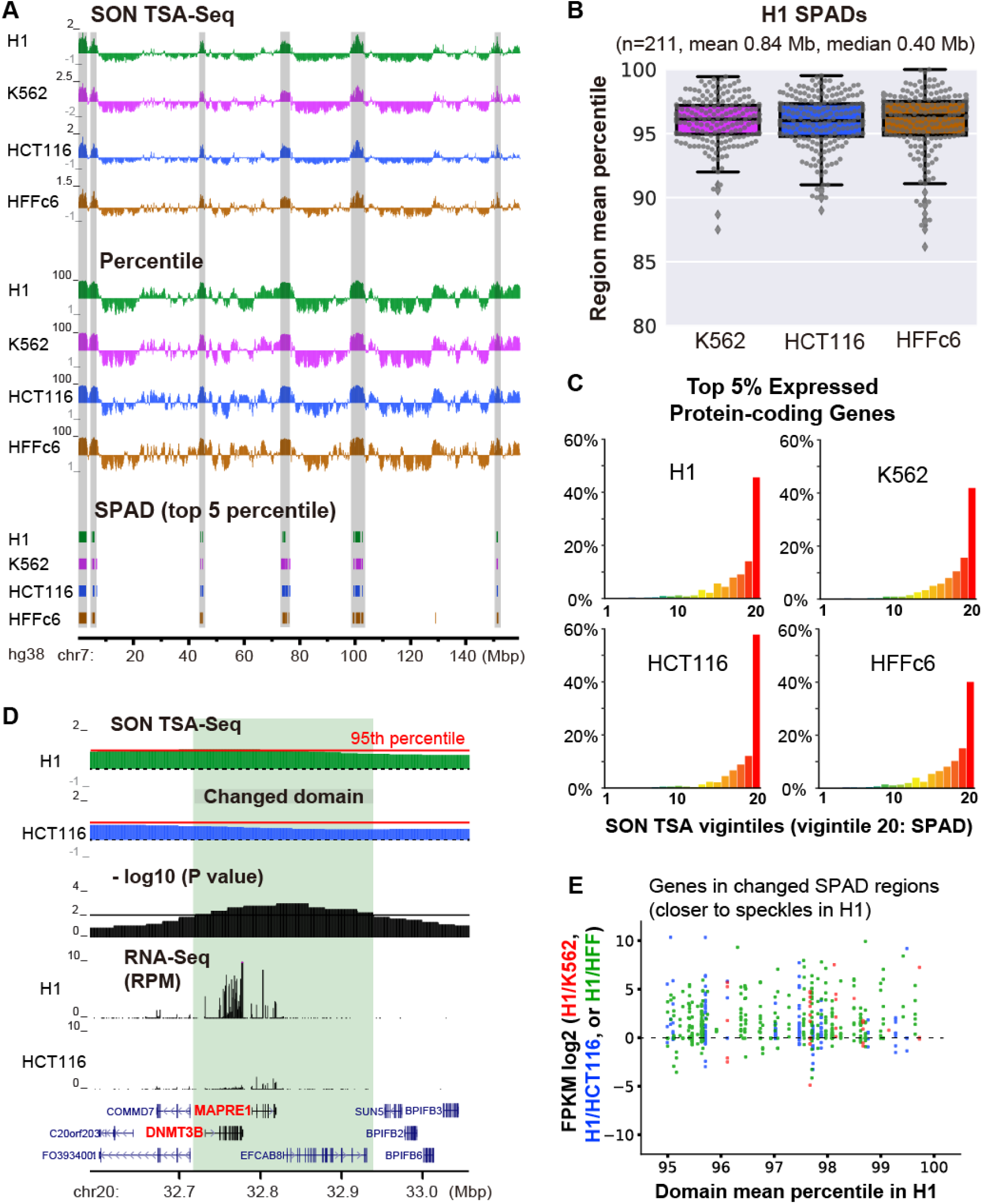
SPADs remain close to nuclear speckles and show high gene expression in all cell lines, yet small position shifts even closer to nuclear speckles correlate with increased gene expression. **A)** SON TSA-Seq enrichment score (top) and SON TSA-Seq signal as a genome-wide percentile score (middle) tracks (20 kb bins), and segmented SPeckle Associated Domains (SPADs) (bottom) in 4 human cell lines (chr7). SPADs were defined as contiguous bins, each with >95th percentile score (Methods). **B)** TSA-Seq score percentile distributions of H1 SPADs in other 3 cell lines. Box plots (with data points as dots) show median (inside line), 25th (box bottom) and 75th (box top) percentiles, 75th percentile to highest value within 1.5-fold of box height (top whisker), 25th percentile to lowest value within 1.5-fold of box height (bottom whisker), and outliers (diamonds). **C)** Distributions of percentages (y-axis) of top 5% expressed protein-coding genes versus SON TSA-Seq vigintiles (division of percentile scores, 0-100, into 20 bins of 5% size each) (x-axis) in 4 cell lines. **D)** Top-to-bottom (light green highlight shows domain repositioned relative to speckles): comparison of H1 and HCT116 smoothed SON TSA-Seq enrichment score tracks (black dashed lines-zero value, red solid lines indicate 95^th^ percentiles of TSA-Seq enrichment scores), -log10 p-values per bin (significance of change in rescaled SON TSA-Seq scores), RNA-Seq RPM values, and gene annotation (GENCODE). **E)** Expression changes of protein-coding genes in SPADs (in H1) closer to speckles in H1: scatterplots show log2 fold-changes in FPKM ratios (y-axis) of H1/K562, H1/HCT116, or H1/HFF versus H1 TSA-Seq score percentiles (x-axis).

Overlapping of SPADs across all 4 cell lines revealed 56.4% (129.3 Mbp) are classified as SPADs (>95^th^ percentile) in all 4 cell lines, 12.5% (28.6 Mbp) in 3 cell lines, 13.0% in 2 cell lines (29.7 Mbp), and 18.1% (41.5 Mbp) in just 1 cell line, out of 229.1 Mbp total (Fig. S5F). Moreover, the majority of SPADs remain near speckles (>90^th^ percentile, and 100% of SPADs >80^th^ percentile in relative SON TSA-Seq enrichment scores) in all 4 cell lines (Fig. 2A-B, Fig. S5A-C), suggesting SPADs remained largely conserved among all 4 cell types, particularly considering the ∼5 percentile differences between biological replicates for regions with top 10% TSA-Seq scores (Fig. S5D-E).

To correlate SPADs with gene expression, we divided human protein-coding genes into 20 groups according to their SON TSA-Seq scores in each cell line. SPADs identified in each cell line were dramatically enriched in the top 5% expressed genes (Fig. 2C) and had the highest FPKM values in all 4 cell lines (Fig. S5G), suggesting SPADs were correlated with genomic domains with constitutively high, overall gene expression-even if individual genes within these domains changed their expression in different cell lines. Also, SPADs were enriched in housekeeping genes compared to non-housekeeping genes (Fig. S5H).

We then asked if statistically significant changes in the speckle proximity of SPADs correlated with changes in gene expression using pair-wise comparisons between cell lines. Starting with all genomic regions defined in at least one of these cell lines as SPADs, we identified subregions 100 kb or larger which showed changed TSA-Seq scores with a p-value threshold of 0.01 in all 20 kb bins within these subregions. These “changed” subregions either were SPADs in both cells (>95^th^ genomic percentile in speckle proximity) that moved even closer to nuclear speckles in one cell line, or were defined as non-SPADs in one cell line but then showed a statistically significant change which resulted in their being classified as SPADs in the other cell line. Changed SPAD regions that were significantly closer to speckles in H1 compared with HFFc6 (HCT116, or K562) accounted for 13.4% (5.2%, or 1.8%) of H1 SPADs, with changes ranging from 1.3 to 13.2 (1.6 to 7.2, or 2.7 to 10.0) and mean changes of 4.4 (4.1, or 5.0) in genomic percentiles measuring proximity to speckles. The p-value threshold of 0.01 was quite stringent considering we were requiring 5 adjacent 20 kb bins, each with a p-value less than this threshold, to call regions that showed significant changes in speckle position.

Strikingly, these small relative changes in SPAD distances to nuclear speckles were accompanied by a pronounced bias towards increased expression of those genes within these SPADs that positioned slightly closer to nuclear speckles in H1 cells (Fig. 2D-E).

### Moderate distance changes relative to nuclear speckles correlate genome-wide with changes in gene expression and histone modifications

We next extended our statistical analysis to look at the correlation between changes in gene position relative to nuclear speckles and changes in gene expression across the entire genome, identifying all domains genome-wide which change in their position in pair-wise comparisons between cell lines. Comparing H1 and HFFc6, we identified 494 domains positioned closer to speckles in HFFc6 (408 kb average size) and 367 domains (396 kb average size) closer to speckles in H1 (e.g. Fig. 3A, highlighted). When comparing replicates, we only detected 2 changed domains using the same threshold, giving a false discovery rate of 0.23%. Similarly, we identified changed regions in H1 vs HCT116 and H1 vs K562 comparisons (Fig. S6A, D).

**Figure 3.**
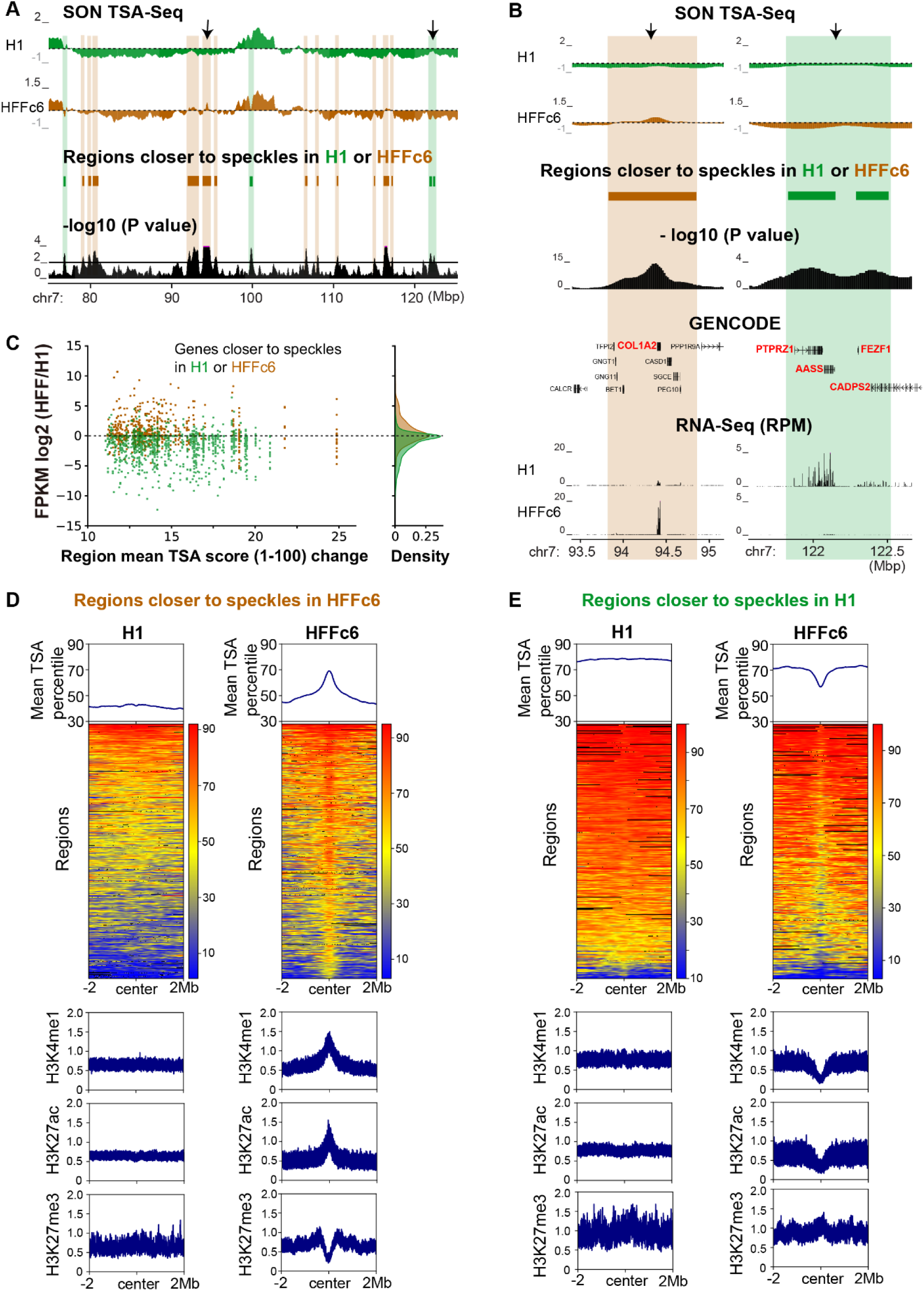
Chromosome regions changing position relative to speckles in H1 versus HFFc6 cells show changes in gene expression and histone modifications. **A)** Top-to-bottom (domains shifting position highlighted): H1 (green) and HFFc6 (brown) smoothed SON TSA-Seq enrichment scores (dashed lines-zero value), domains repositioned relative to speckles, and -log10 p-values per bin (significance of change in rescaled SON TSA-Seq scores). **B-C)** Strong bias towards increased gene expression in domains shifting closer to nuclear speckles: **B)** Zoomed view of two regions (arrows in **A)** plus gene annotation (GENCODE) and RNA-Seq RPM values. **C)** Scatterplots show log2 fold-changes in FPKM ratios (y-axis) between HFFc6 and H1 versus absolute values of changes in mean scaled TSA-Seq scores (x-axis). Green (brown) dots are genes closer to speckles in H1 (HFFc6). Kernel density plots show gene density (right). **D-E)** TSA-Seq percentiles and histone modifications fold-enrichment for regions significantly closer to speckles in HFFc6 (**D**) or H1 (**E**) +/- 2 Mbps from each region center. Top: TSA-Seq percentiles averaged over all regions; Middle: heatmaps with color-coded TSA-Seq score percentiles; Bottom: histone modification fold-enrichment for H3K4me1, H3K27Ac or H3K27me3 averaged over all regions.

Surprisingly, we detected no changed domains that showed large shifts in SON TSA-Seq scores. To compare TSA-Seq scores from different cell lines we first normalized these scores on a scale from 1 to 100, where 1 corresponds to the SON TSA-Seq score estimated minimum and 100 to the SON TSA-Seq score estimated maximum (See Supplementary Methods). On this normalized scale, statistically significant observed changes ranged from ∼10 – 26, representing relatively modest shifts in nuclear position. We used our distance calibration in K562 cells (Supplementary Table 1 and 2, Methods) to convert SON TSA-Seq score into estimated mean distance to nuclear speckles. Estimated distances to speckles in K562 cells were linearly correlated with the scaled TSA-Seq scores (1-100) that we used for the statistical pair-wise comparison (Fig. S7A). Based on this conversion, domains that significantly shifted in their relative nuclear speckle distance in the comparison of H1 and K562 cells, showed a predicted mean distance shift ranging from 0.113 to 0.375 µm relative to the distance to nuclear speckles expected if there had not been a change in their TSA-Seq score (Fig. S7B).

Despite the relatively modest magnitude of these position shifts relative to nuclear speckles, again we saw a correlation with changes in gene expression, even with the vast majority of changed domains corresponding to non-SPAD domains. Chromosome regions closer to nuclear speckles in one cell line versus the other typically showed one or more genes with higher relative gene expression levels in this cell line (e.g. Fig. 3B, Fig. S6B, E), including many examples of genes with tissue-specific expression-for example, Collagen genes in fibroblasts (Fig. 3B, Fig. S8). Genome-wide analysis confirmed this marked expression bias (Fig. 3C, Fig. S6C, F). In the comparison of H1 vs HFFc6 cells, of 628 genes with log2-fold changes >=1, 379 genes >= 2, and 248 genes >= 3 located within chromatin domains closer to speckles in H1 versus HFFc6, 91%, 94%, and 96% of these genes, respectively, show higher expression in H1. Of 307 genes with log2-fold changes >= 1, 161 genes >=2, and 94 genes >=3 located within domains closer to speckles in HFFc6 versus H1, 66%, 77%, and 83% of these genes, respectively, show higher expression in HFF.

Examining all changed domains in the H1 versus HFFc6 pair-wise comparison, revealed an interesting asymmetry in the typical pattern of relative nuclear speckle distance shifts (Fig. 3D, E). Regions that shifted closer to nuclear speckles in HFFc6 relative to H1 cells, showed relatively flat, plateau TSA-Seq score profiles in H1 cells changing to local peaks in SON TSA-Seq scores in HFFc6 cells (Fig 3D, top); these local peaks were ∼600 kb in full width at half-maximum (FWHM), representing a localized movement towards nuclear speckles. These local peaks were associated with changes in histone modifications, with increased active and decreased repressive histone marks correlated with regions moving closer to speckles (Fig. 3D bottom). Conversely, regions that shifted relatively further from nuclear speckles in HFFc6 cells showed relatively flat, plateau TSA-Seq score profiles in H1 cells changing to local valleys in SON TSA-Seq scores in HFFc6 cells (Fig 3E, top). These valleys had ∼400 kb FWHM and showed decreased active and increased repressive histone marks (Fig. 3E, bottom). This asymmetry suggests localized, directed movements of ∼400 – 600 kb domains towards or away from nuclear speckles as a result of cell differentiation that highly correlate with gene up- or down-regulation, respectively. These chromosome domains moving towards speckles in HFFc6 cells, generally showed low SON TSA-Seq scores in H1 versus intermediate scores in HFFc6 (Fig. 4A, scores averaged over domains). Conversely, chromosome domains moving away from speckles in HFFc6 cells, showed intermediate to high SON TSA-Seq scores in H1 cells (Fig. 4B, scores averaged over domains).

**Figure 4.**
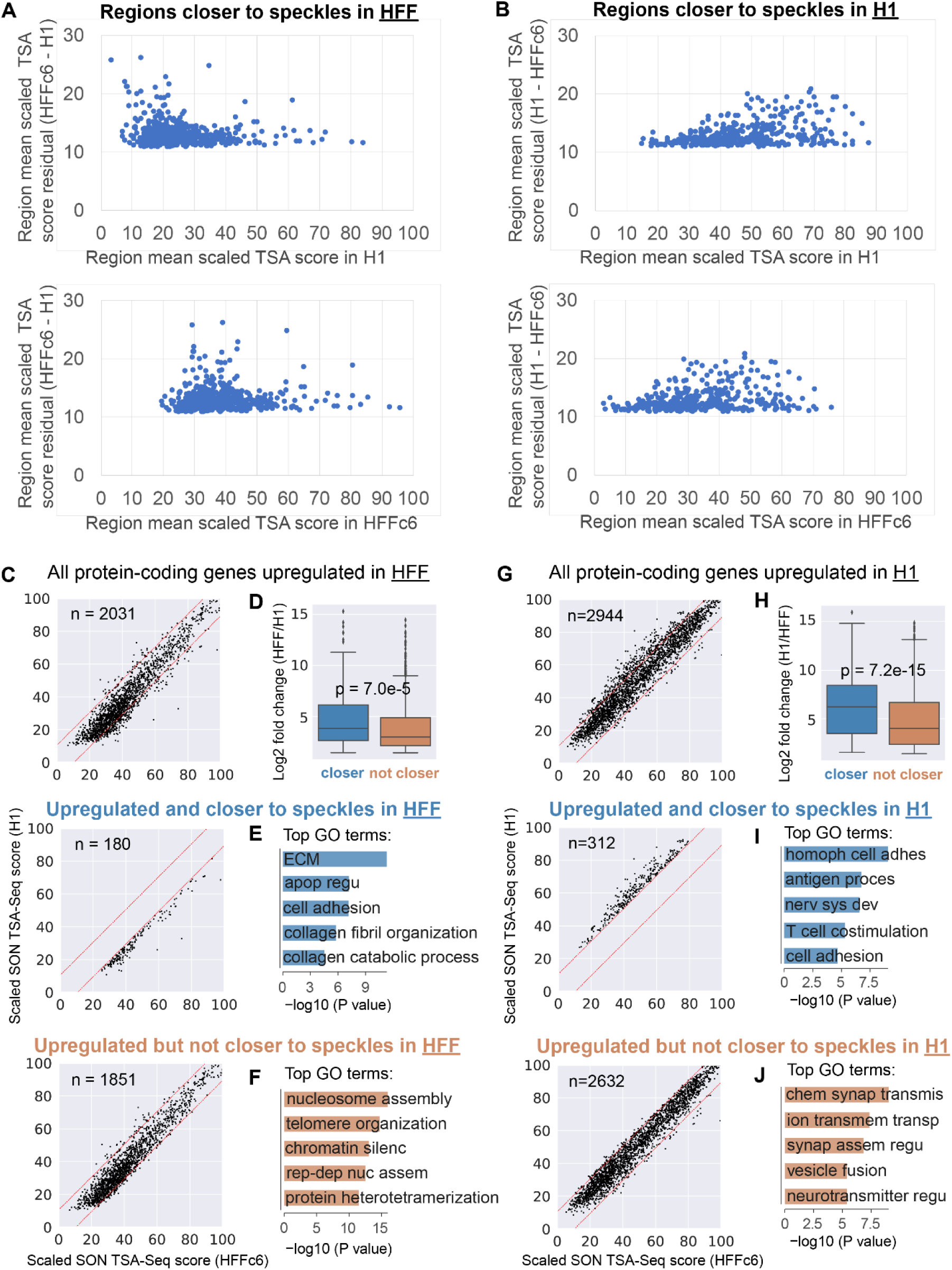
Shifts in domain positioning relative to speckles are small and correlate with changes in expression of cell-type specific genes. **A, B)** Scatter plots show change in scaled SON TSA-Seq scores (1-100 range) averaged over each domain (y-axis) with a significant score change for domains closer to speckles in HFFc6 (A) or H1 (B) as function of scaled TSA-Seq scores averaged over each domain (x-axis) in H1 (top) or HFFc6 (bottom). **C, G)** Scatter plots show scaled SON TSA-Seq scores (1-100 range) in H1 hESCs (y-axis) versus HFFc6 (x-axis) for protein-coding genes (dots) that are upregulated in HFF (**C**) or in H1 (**G**): top panels-all upregulated genes, middle panels-upregulated genes in regions closer to nuclear speckles in HFFc6 (**C**) or in H1 (**G**), bottom panels-upregulated genes in regions that are not closer to nuclear speckles. Red dashed lines represent thresholds for statistically significant changes in scaled HFFc6 vs H1 SON TSA-Seq scores. **D, H)** Comparison of log2-fold changes in gene expression for protein-coding genes significantly upregulated in HFF (**D**, HFF/ H1) or in H1 (**H**, H1/HFF) for: upregulated genes in regions closer to nuclear speckles in HFFc6 (**D**, left, blue) or in H1 (**H**, left, blue) versus upregulated genes in regions that are not closer to nuclear speckles in HFFc6 (**D**, right, orange) or in H1 (**H**, right, orange) (Welch’s t-test). Box plots show median (inside line), 25^th^ (box bottom) and 75^th^ (box top) percentiles, 75^th^ percentile to highest value within 1.5-fold of box height (top whisker), and 25^th^ percentile to lowest value within 1.5-fold of box height (bottom whisker). **E, F, I, J)** Top gene ontology (GO) terms for protein-coding genes upregulated in HFF (**E, F**) or in H1 (**I, J**) for upregulated genes in regions closer to nuclear speckles in HFFc6 (**E**) or in H1 (**I**) versus upregulated genes in regions not closer to nuclear speckles in HFFc6 (**F**) or in H1 (**J**). Abbreviated GO terms: **E)** ECM: extracellular matrix organization; apop regu: negative regulation of apoptotic process; **F)** chromatin silenc: chromatin silencing at rDNA; rep-dep nuc assem: DNA replication-dependent nucleosome assembly; **I)** homoph cell adhes: homophilic cell adhesion via plasma membrane adhesion molecules; antigen proces: antigen processing and presentation of peptide or polysaccharide antigen via MHC class II; nerv sys dev: nervous system development; **J)** chem synap transmis: chemical synaptic transmission; ion transmem transp: ion transmembrane transport; synap assem regu: positive regulation of synapse assembly; neurotransmitter regu: calcium ion-regulated exocytosis of neurotransmitter.

### Genes with significantly higher expression in regions that position closer to nuclear speckles are enriched in cell-type specific functions

The previous section described the pronounced bias for genes in regions that shift closer to nuclear speckles to show increased rather than decreased gene expression in that cell line. If we instead ask how many differentially expressed (DE) genes in two cell lines are in chromosome regions that change their position significantly relative to nuclear speckles, we find ∼9-fold more genes with higher gene expression in one cell line versus another that did not position closer relative to nuclear speckles in that cell line versus regions that did position closer to nuclear speckles in the given cell line (Fig. 4C, G). The difference between DE genes in regions that positioned closer to nuclear speckles in a given cell line versus DE genes in regions that did not position closer did not appear related to the magnitude of changes in gene expression. DE genes located in chromatin domains that moved closer to nuclear speckles showed a moderate shift in their distribution towards higher fold-changes in expression, although there was extensive overlap in expression changes with DE genes in chromatin domains that did not move closer (Fig. 4D, H).

Instead, gene ontology (GO) analysis suggests DE genes in regions closer to speckles in one cell line are enriched in genes with cell type specific functions, including extracellular matrix organization and collagen related functions for regions closer to speckles in HFFc6 fibroblasts (Fig. 4E) and homophilic cell adhesion related to stem cell colony formation and maintenance (Pieters and van Roy 2014) in H1 hESCs (Fig. 4I). “Cell adhesion” function was enriched in DE genes that were closer to speckles in either H1 or HFFc6 cells (Fig. 4E, I), suggesting the same functional but different groups of genes were regulated by closer localization to nuclear speckles in the two different cell types.

### Genome-wide analysis of changes in genome organization relative to nuclear speckles after heat-shock reveals partial “hardwiring” of heat-shock protein gene loci

To further investigate the possible “hardwiring” and dynamics of genome organization relative to nuclear speckles, we applied TSA-Seq mapping to K562 cells undergoing heat shock. As previously reported (Spector and Lamond 2011), nuclear speckles rounded up and enlarged upon 30 min, 1h, or 2h heat shock at 42°C (Fig. SF9A). Live-cell imaging recently has revealed that at least some of this speckle enlargement after heat shock occurs through directed movement of small speckles towards larger nuclear speckles, followed by speckle fusion (Kim et al. 2019). Surprisingly, despite these significant changes in nuclear speckle size, shape, and movement and a dramatic gene expression profile change, including thousands of genes down-regulated upon 30 min heat shock (42°C) (Vihervaara et al. 2017), we found nearly no significant genome-wide changes in SON TSA-Seq scores after 30 min to 2 hrs heat shock (Fig. S9B). Differences in TSA-Seq scores between control and heat shocked samples were largely comparable to differences between biological replicates (Fig. S9C).

These unchanged chromosome regions included 4 of 8 chromosome loci-*HSPA1A/HSPA1B/HSPA1L, DNAJB1, HSP90AA1, and HSP90AB1* (Fig. 5A-B, Fig. S10)-containing major induced heat shock protein (HSP) genes (based on PRO-Seq in this same K562 cell line (Vihervaara et al. 2017)). Interestingly, each of these four heat shock gene loci are positioned both before and after heat shock very near nuclear speckles, scoring near the 100^th^ genomic percentile in speckle proximity. This is striking, given our recent demonstration of increased (decreased) nascent *HSPA1A* transcript production within several minutes after nuclear speckle association (disassociation) of a *HSPA1B* BAC transgene and a several-fold increased gene expression of the endogenous HSPA1A/HSPA1B/HSPA1L locus in human HAP1 cells (Kim et al. 2020). Indeed, the *HSPA1A/HSPA1B/HSPA1L* endogenous chromosome locus is in the 100^th^ percentile in speckle proximity in K562 cells even before heat shock, and thus appears pre-positioned in nearly all cells adjacent to nuclear speckles where its gene expression is induced to maximal levels (Kim et al. 2020).

**Figure 5.**
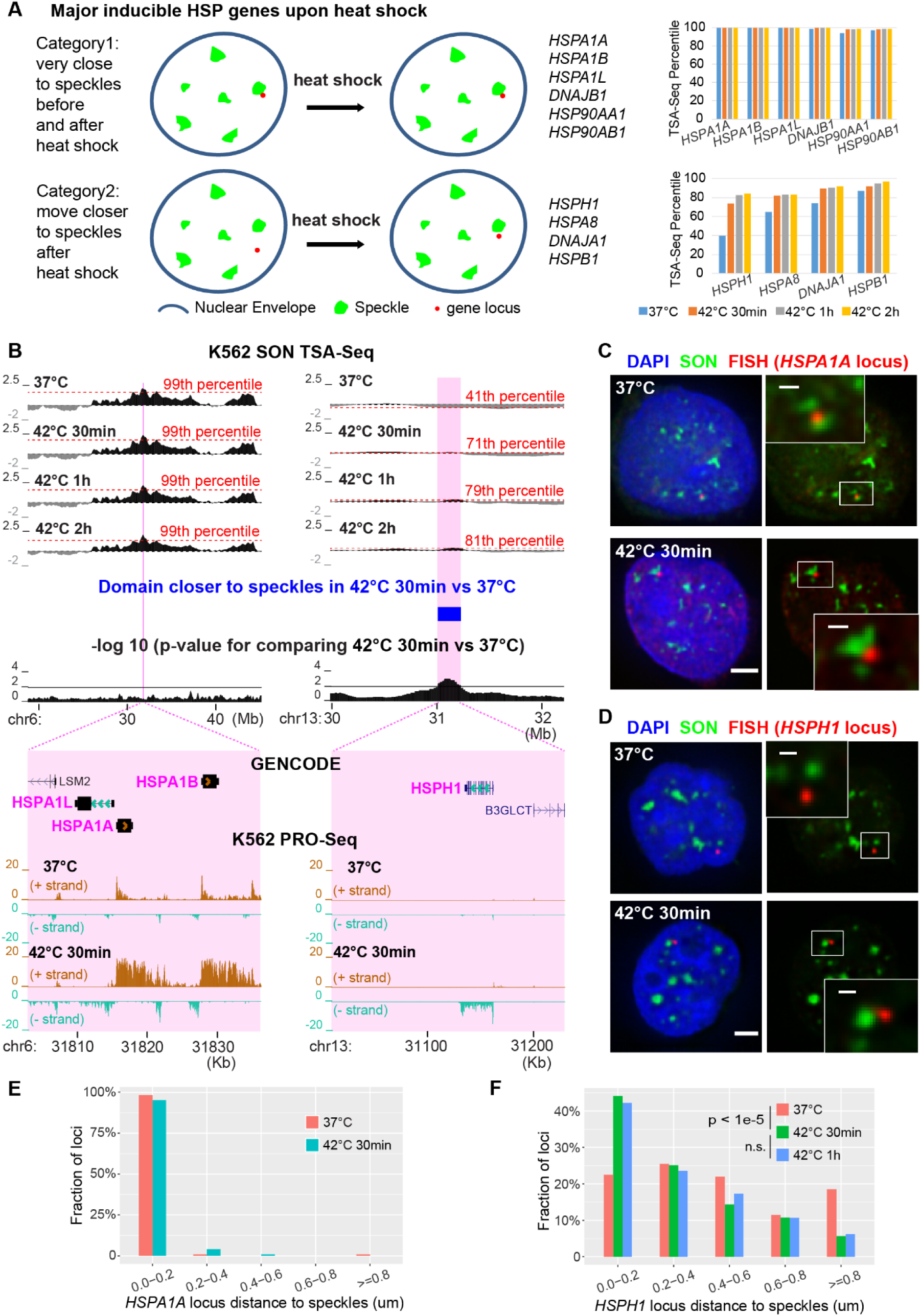
Approximately half of inducible heat shock protein gene loci are pre-positioned very close to nuclear speckles in SPADs while remaining gene loci shift closer to speckles after heat shock. **A)** Schema (left) and TSA-Seq percentiles at 0 (blue), 30 mins (orange), 1 hr (grey), and 2 hrs (gold) after heat shock (right) for genes pre-positioned near speckles (top) versus those moving closer after heat shock (bottom). **B)** TSA-Seq and PRO-Seq profiles of *HSPA1A* (left) and *HSPH1* (right) loci in K562 cells after heat shock. Top to bottom: smoothed SON TSA-Seq enrichment scores for control or heat shock times (red dashed lines-labeled TSA-Seq score percentiles), significantly changed domains comparing 37 °C vs 42 °C 30 mins (blue, right), - log10 p-values for comparison of rescaled SON TSA-Seq scores between 37 °C and 30 mins heat shock (20 kb bins), and zoomed view of the highlighted two regions (magenta) plus gene annotation (GENCODE) and K562 PRO-Seq tracks showing normalized read count (BPM, 50 bp bins). **C-D)** 3D immuno-FISH for *HSPA1A* (red, **C**) or *HSPH1* (red, **D**) plus SON immunostaining (green) and merged channels with DAPI (blue, left, scale bars: 2 μm) in K562 cells at 37 °C (top) or at 42 °C for 30 mins (bottom). Insets: 3× enlargement of the white-boxed image area (right, scale bars: 0.5 μm). **E-F)** Nuclear speckle distance distributions for *HSPA1A* (**E**) or *HSPH1* (**F**) FISH signals in control versus after heat shock. *HSPA1A*: n=119 (37 °C) or 124 (42 °C 30 mins). *HSPH1*: n=200 (37 °C) or 195 (42 °C 30 mins) or 225 (42 °C 1 hr), combination of two biological replicates; Chi-Square Test of Homogeneity, p = 3.793 × 10^−6^ between 37 °C and 42 °C 30 mins, p = 0.9341 (n.s.) between 42 °C 30 mins and 42 °C 1h.

We did identify 13 chromosome regions, ∼100 – 680 Kbp in size and together comprising 3.36 Mbp, positioned closer to speckles after 30 mins heat shock. Four of these 13 regions corresponded to the remaining 4 of 8 heat shock protein gene chromosome loci containing major induced HSP genes as identified by PRO-Seq (Vihervaara et al. 2017): *HSPH1, HSPA8, DNAJA1, HSPB1* (Fig. 5A-B, Fig. S10). Their position shifts closer to nuclear speckles with heat shock were modest, in the same range as observed previously for regions that changed relative speckle position in pair-wise comparisons of cell lines (Fig. 3C, Fig. S6C, F). Specifically, they ranged from ∼5 – 30 percentile changes in genomic percentiles measuring proximity to speckles (30, *HSPH1* locus; 17, *HSPA8* locus; 15, *DNAJA1* locus; 5, *HSPB1* locus) and ∼11 – 13 scaled (1-100) SON TSA-Seq score changes (13, *HSPH1* locus; 12, *HSPA8* locus; 12, *DNAJA1* locus; 11, *HSPB1* locus).

### *HSPA1A/HSPA1B/HSPA1L* deterministic positioning near nuclear speckles; *HSPH1* movement towards nuclear speckles and large-scale chromatin decondensation after heat shock associated with increased nascent transcripts

We next applied 3D immuno-FISH to investigate the nuclear positioning of induced HSP genes relative to nuclear speckles. We used the *HSPA1A/HSPA1B/HSPA1L* locus as an example of a heat-shock locus positioned near nuclear speckles both before and after heat shock and the *HSPH1* locus as an example of a heat-shock locus that moved closer to nuclear speckles after heat shock.

DNA FISH showed near deterministic positioning of the *HSPA1A/HSPA1B/HSPA1L* locus near nuclear speckles in K562 cells. Approximately 90% – 100% alleles localized within 0.2 µm from speckles before and after heat shock (0.024 µm and 0.040 µm mean, 0 µm and 0 µm median distances before and after 30 mins heat shock) (Fig. 5C, E, Fig. S11A). Similar deterministic positioning of this locus near speckles was seen in four additional human cell lines without or with heat shock: WI38 and IMR90 normal fibroblasts, hTERT-immortalized Tig3 fibroblasts, and HCt116 cells (Fig. S11B-E).

In contrast, the *HSPH1* locus showed a significant shift in the observed distribution of distances from nuclear speckles at 30 mins after heat shock which was maintained at 1 hr after heat shock (Fig. 5D, F). In particular, there was an ∼2-fold increase in alleles localized within 0.2 μm from nuclear speckles at 30 min and 1 hr after heat shock, from ∼20% to ∼40% of alleles, together with an ∼4-fold reduction in alleles localized 0.8 μm or further from nuclear speckles, from ∼20% to ∼5% of alleles (Fig. 5F).

Interestingly, in addition to the shift in *HSPH1* position towards nuclear speckles, we also observed an ∼3-fold increase (from ∼20% to ∼60%) in *HSPH1* alleles showing a decondensed DNA FISH signal (“dec”, Fig. 6A-B) larger than the microscopy diffraction limit in size, typically observed for *HSPH1* alleles associated with nuclear speckles after heat shock (Fig. 6A-B). These decondensed alleles appeared as elongated, fiber-like FISH signals, frequently with one region of the FISH signal touching a nuclear speckle (Fig. 6A). In contrast, FISH signals that appeared as round, diffraction-limited spots (“non-dec”), represented ∼80% of *HSPH1* alleles before heat shock. Decondensed alleles showed an overall shift in their distance distribution closer to nuclear speckles as compared to non-decondensed alleles, with a >2-fold increase (from ∼20% to ∼55%) for alleles localized within 0.2 μm from nuclear speckles (Fig. 6C).

**Figure 6.**
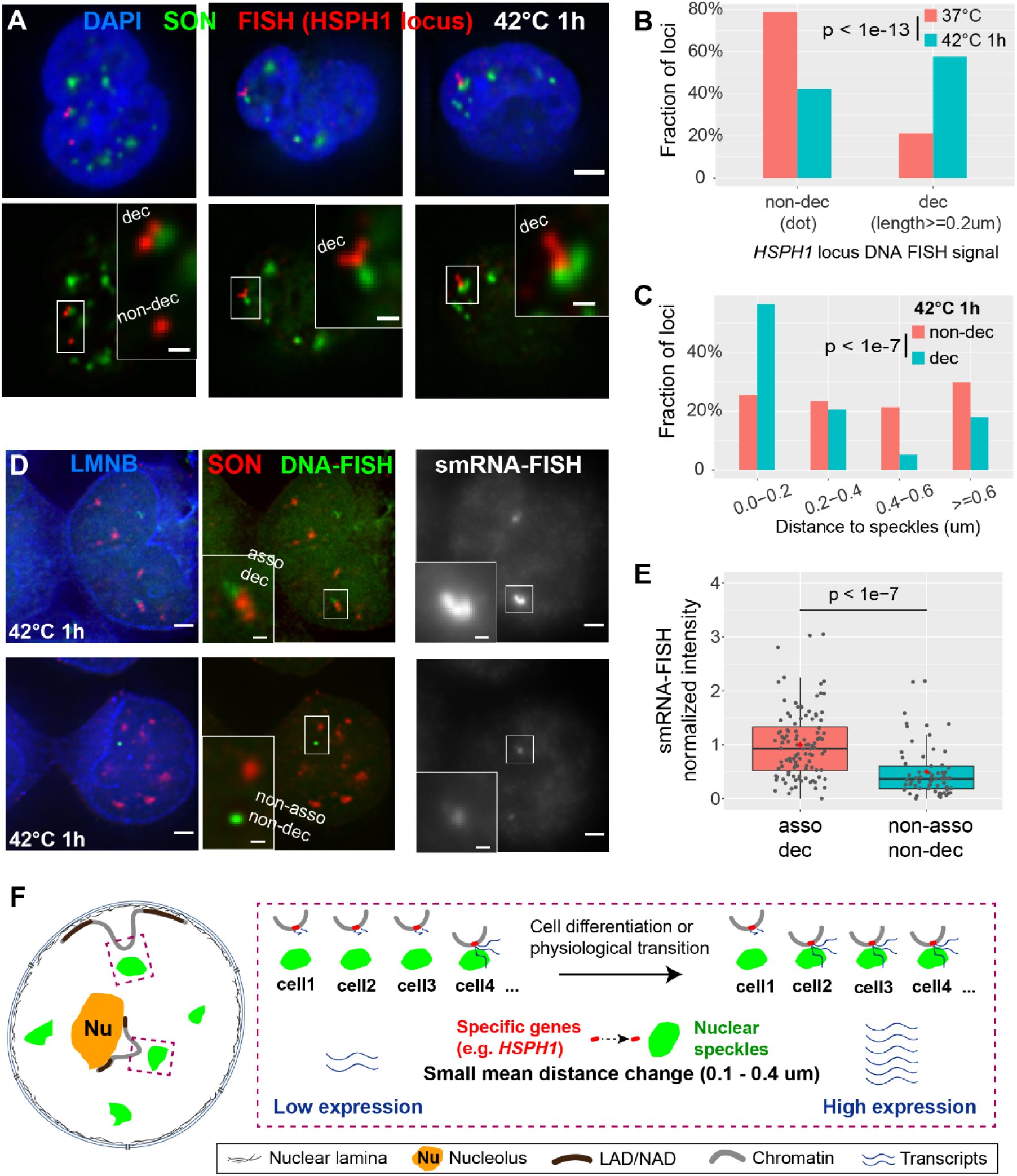
Both distance shift towards speckles and large-scale chromatin decondensation of *HSPH1* locus after heat shock correlate with gene expression amplification in K562 cells. **A)** 3D immuno-FISH for *HSPH1* locus (red) plus SON immunostaining (green) and merged channels with DAPI (blue, top, scale bar: 2 μm) in K562 cells after heat shock (42 °C 1h). Types of FISH signals-left: decondensed (upper, elongated) and non-decondensed signals (lower, dot); middle and right: decondensed signals. Insets: 3× enlargement of region in white-box (bottom, scale bars: 0.5 μm). **B)** Increase in decondensed *HSPH1* loci after heat shock: n=198 (37 °C) or 224 (42 °C 1 hr), combination of two biological replicates; Chi-Square Test of Homogeneity, p = 3.045 × 10^−14^. **C)** Speckle distance distributions for different cases of *HSPH1* locus FISH signals in K562 cells after 1 hr heat shock (42 °C): n=141 (non-decondensed, dot) or 200 (decondensed, >= 0.2 μm in length), combination of two biological replicates; Chi-Square Test of Homogeneity, p = 2.74 × 10^−8^ between non-decondensed and decondensed. **D)** Combined DNA- and single-molecule RNA (smRNA)- immuno-FISH for *HSPH1* locus showing DNA-FISH signal (green), SON immunostaining (red), Lamin B immunostaining signal (blue), and smRNA-FISH (grey) in K562 cells after 1 hr heat shock (scale bars: 2 μm). Decondensed chromatin structure (elongated DNA-FISH signal) with speckle association (top) and non-decondensed (dot DNA-FISH signal) not associated with speckles (bottom) and their corresponding smRNA-FISH signal (right). Insets: 3× enlargement of white-boxed area (scale bar: 0.5 μm). Single optical section is shown containing center (in z) of DNA FISH signal ; RNA-FISH images correspond to projected sum of 20 optical sections. **E)** Normalized smRNA-FISH signals corresponding to different chromatin states based on DNA immuno-FISH. Combination of two biological replicates-asso dec: distance to speckles < 0.4 µm and signal length >= 0.4 µm, n=108, mean=1.00, median=0.93; non-asso non-dec: distance >= 0.6 µm and dot signal, n=68, mean=0.50, median=0.37. Box plots (data points-grey dots, means-red dots) show median (inside line), 25th (box bottom) and 75th (box top) percentiles, 75th percentile to highest value within 1.5-fold of box height (top whisker), 25th percentile to lowest value within 1.5-fold of box height (bottom whisker). Welch’s t-test: p = 1.179 × 10^−8^. **F)** Cartoon model: small mean distance changes relative to nuclear speckles predicted by TSA-Seq may reflect changes in the distribution of distances such that an increased fraction of alleles shows close association with nuclear speckles with an accompanying amplification of gene expression.

We combined single-molecule RNA FISH (smRNAFISH) with subsequent DNA FISH to correlate differential *HSPH1* expression with both position relative to nuclear speckles and large-scale chromatin decondensation of the locus. Based on DNA-FISH, we defined two opposite subclasses of alleles: “speckle-associated with decondensation” (distance to speckles < 0.4 µm and signal length >= 0.4 µm) or “non-speckle-associated without decondensation” (distance to speckles >= 0.6 µm and dot signal). Our rationale was that non-decondensed, speckle-associated alleles or non-speckle associated, decondensed alleles might represent intermediate, transient states. We used a smRNA FISH probe set spanning the 5’ UTR (2 oligos) and 1^st^ intron (28 oligos) of the *HSPH1* gene to measure nascent transcript levels. Speckle-associated, decondensed *HSPH1* alleles show an ∼ 2.0-fold mean (2.5-fold median) increased nascent transcript smRNA FISH signal relative to non-speckle associated, non-decondensed alleles (Fig. 6D-E,). This 2-fold-increase is in the range of increased nascent transcript signals with speckle-association recently observed for the *HSPA1A/HSPA1B/HSPA1L* endogenous locus and nearby flanking genes (Kim et al. 2020), suggesting that movement of the *HSPH1* locus to nuclear speckles may similarly be associated with amplification of gene expression.

## Discussion

Our results suggest a surprisingly conserved genome organization relative to nuclear speckles. Approximately, 84% of the genome showed no significant position change relative to nuclear speckles in comparisons of hESCs with all three differentiated cell types – fibroblasts (HFFc6), eythroleukemia (K562) and colon carcinoma (HCT116) – and ∼90% unchanged in single, pair-wise comparisons with hESCs. Meanwhile, the small percentage of the genome that did change position showed only modest estimated mean distance changes relative to nuclear speckles. Yet these modest position changes, concentrated over localized chromatin domains ∼100 Kbp – 5.8 Mbp in size, highly correlated with changes in histone marks linked to gene activity and actual gene expression changes; shifts towards (away from) speckles were biased strongly towards increased (decreased) expression, particularly for cell-type specific genes. This conclusion of largely “hardwired” genome organization relative to nuclear speckles was strengthened by the observation of prepositioning of 4/8 of the major heat shock induced loci very close (top 1-2% of the genome) to nuclear speckles prior to heat shock. The remaining 4 major heat-shock induced loci were a significant fraction of the 13 genomic loci (∼0.1% of genome) that moved closer to nuclear speckles within 30 mins of heat shock. Like the *HSPA1A* locus (Kim et al. 2020), the movement and then nuclear speckle association of the HSPH1 locus correlated with a higher expression of nascent transcripts.

These results are surprising based on previous literature. A series of papers in the late 1990s and early 2000s from the Lawrence laboratory demonstrated the preferential localization near nuclear speckles of approximately half of about 25 highly expressed genes (Hall et al. 2006). Several of these speckle-associated genes, as well as others described in the literature, were developmentally regulated genes expressed at high levels in specific differentiated cell types, including collagen 1A1 in fibroblasts (Smith et al. 1999), cardiac myosin heavy chain (MyHC) in myotubes (Smith et al. 1999), mono-allelically expressed GFAP in astrocytes (Takizawa et al. 2008a), and beta-globin in erythroblasts (Brown et al. 2006). Two of these developmentally-regulated genes localized away from nuclear speckles in cell types in which these genes were inactive: the MyHC gene localized near the nucleolus in fibroblasts (Xing et al. 1995), while the human beta-globin gene localized near the nuclear periphery in pre-erythrocytes (Brown et al. 2006) and fibroblasts (Bian et al. 2013). Thus, it was natural to expect significant numbers of genes showing large movements relative to nuclear speckles. This expectation was further increased by the direct visualization by live-cell imaging of directed movements up to 5 microns in length from the nuclear periphery to nuclear speckles of *HSPA1A* plasmid transgene arrays after heat-shock induced transcriptional (Khanna et al. 2014).

More recently, two results led us to question these expectations of large gene movements relative to nuclear speckles. First, was our discovery that Type I transcription hot-zones, closest to nuclear speckles, showed larger genomic distances to the nearest LAD boundary than Type II transcription hot-zones, at intermediate distance to nuclear speckles. We speculated that the larger distance from the nearest LAD for Type I hot-zones might have evolved to facilitate contact with nuclear speckles which are relatively depleted from the nuclear periphery. Second, we realized that in contrast to heterochromatic, multi-copy plasmid *HSPA1A* transgene arrays, which frequently localized near the nuclear periphery, Hsp70 BAC transgenes instead appeared to show a very high fraction of alleles localizing at nuclear speckles after random integration into CHO cells, similar to the ∼90% localization to nuclear speckles of the endogenous locus in the same CHO cells (Hu et al. 2009; Kim et al. 2020).

We now show the *HSPA1A/HSPA1B/HSPA1L* locus and 3 other major heat shock loci are even more strongly associated with nuclear speckles in all 5 human cell lines examined here than the same locus in CHO cells. Notably, live-cell imaging recently revealed that HSPA1B transgenes not already located near nuclear speckles required an additional several minutes to move to nuclear speckles and reach maximum nascent transcript production (Kim et al. 2020). Thus, this “pre-wiring” of the endogenous *HSPA1A/HSPA1B/HSPA1L* locus very close to nuclear speckles likely ensures a robust heat shock activation across all alleles in all cells. We anticipate a similar effect of close speckle proximity on the robust activation of the other three heat shock gene loci, and possibly other stress-induced genes, that are pre-wired near nuclear speckles. More generally, our observed positioning of SPADs, defined in one cell type, close to nuclear speckles in other cell lines suggests that this pre-wiring concept may apply to a much larger set of genes.

Likewise, it is striking that the modest shift towards nuclear speckles of the remaining 4 major heat shock loci cover much of the similar range for the modest shifts observed in comparison of the different cell lines. Shifts in the position of chromatin domains between pairs of cell types ranged from ∼10-26 in scaled SON TSA-Seq scores (1-100), corresponding to speckle proximity genomic percentile shifts anywhere from a few to ∼50 percentiles and actual estimated mean distance shifts of only 0.113-0.375 microns. Yet these domain shifts were again highly correlated with significant gene expression changes. Our *HSPH1* microscopy results demonstrate how small changes in ensemble TSA-Seq mapping scores may in fact correspond to significant shifts in the fraction of alleles in close contact with nuclear speckles. This fraction of alleles in close contact with nuclear speckles may undergo gene expression amplification, similar to that observed now for both the HSPA1A/HSPA1B/HSPA1L (Kim et al. 2020) and HSPH1 loci (Fig. 6F).

One still puzzling aspect of our results is that this bias in differential gene expression is not only seen for regions that gain or lose very close proximity to nuclear speckles, but also for regions that shift closer to nuclear speckles but without reaching very close proximity. We speculate that the functional significance of these intermediate distance shifts relative to nuclear speckles may instead be related to shifts in their distance relative to other, still unknown nuclear compartments, for instance RNA polII clusters/condensates (Cisse et al. 2013; Boehning et al. 2018; Cho et al. 2018; Guo et al. 2019), which themselves are positioned nonrandomly relative to nuclear speckles.

With the development of our improved, “super-saturation” TSA-Seq method, we are now better positioned to explore how the genome is organized relative to other nuclear compartments, in addition to nuclear speckles. This includes both known compartments, including the nuclear lamina, nuclear pores, PML bodies, Cajal bodies, and pericentric heterochromatin, as well as currently less well-characterized compartments identified by interesting antibody staining patterns. TSA-Seq is particularly well-suited to explore the genome spatial relationship to liquid-liquid phase-separated condensates that might otherwise be difficult to map by methods such as DamID, ChIP-Seq, or CUT&RUN. Whereas the TSA-Seq 1.0 procedure required sequential staining of several batches of cells over 1-2 months, our new TSA-Seq 2.0 procedure requires 10-20-fold fewer cells performed in a single, one-week staining for nuclear speckle mapping, with a proportional reduction in cell culture costs (for hESCs, from ∼$2000 to ∼$200 per replicate).

These future extensions of TSA-Seq to multiple nuclear compartments in many cell types and states should greatly increase our understanding of the role of nuclear compartmentalization in nuclear genome organization and function.

## Methods

### Cell culture

K562 cells were obtained from the ATCC and cultured following the ENCODE Consortium protocol (http://genome.ucsc.edu/ENCODE/protocols/cell/human/K562_protocol.pdf). H1-ESC (WA01), HCT116, HFF-hTert-clone 6 cells were obtained through the 4D Nucleome Consortium and cultured according to 4DN SOPs (https://www.4dnucleome.org/cell-lines.html). A detailed description for reagents and culturing procedures is provided in **Supplementary methods.**

### Coverslip TSA staining & TSA-Seq

TSA staining and TSA-Seq procedures were modified from our previous publication (Chen et al. 2018). TSA labeling conditions are summarized in Fig. S1A. Detailed procedures are provided in **Supplementary methods**, including for TSA staining of cells attached to/grown on coverslips, TSA-Seq for suspension cells (K562), TSA-Seq for attached cells (H1, HFFc6, HCT116), and dot blot estimation of DNA biotinylation levels.

### Microscopy

3D optical sections were acquired with 0.2-μm z-steps using DeltaVision SoftWoRx software (GE Healthcare) and either an Applied Precision Personal DeltaVision microscope system with a 60× oil objective (NA 1.4) and a CoolSNAP HQ2 charge-coupled device camera or a DeltaVision OMX microscope system with a 100× (NA 1.4) objective lens and an Evolve 512 Delta EMCCD Camera operated in wide-field mode. Image deconvolution and registration also were done using SoftWoRx.

Image intensity line profiles were measured with FIJI software (ImageJ, NIH) using the Plot Profile function and normalized by exposure time and the % transmitted exciting light. Normalized intensity plots were generated using OriginPro 2018 software (OriginLab).

### TSA-Seq data processing

We used a similar pipeline as in a previous paper (Chen et al. 2018) to process TSA-Seq data. Briefly, we mapped raw sequencing reads to the human reference genome (hg38, Chromosome Y excluded for female cell line K562) using Bowtie2 (Langmead and Salzberg 2012) (version 2.1.0) with default parameters. We applied the rmdup command from SAMtools (Li et al. 2009) (version 1.5) to remove potential PCR duplicates in the alignment. We then used these alignment files as the input files for TSA-Seq normalization, as described in detail in **Supplementary methods**.

### TSA distance prediction and comparison between different staining conditions

To convert TSA-Seq scores into mean speckle distances, we previously (Chen et al. 2018) fit the TSA-Seq enrichment ratios, y (the TSA-Seq ratio prior to the log2 operation in the TSA-Seq normalization procedure), and mean speckle distances, x (measured by 3D immuno-FISH), corresponding to multiple genomic loci, to the calibration equation 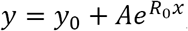, to obtain constants *y*_0_, A and *R*_0_. Here we applied a new “hybrid” method, which instead estimates *R*_0_ by FISH calibration and estimate *y*_0_ and *A* using minimum and maximum TSA-Seq enrichment ratios, as described in **Supplementary methods**.

To compare these speckle distances estimated from TSA-Seq data produced using two different TSA staining conditions, we generated histograms of their distance residuals over each 20 kb bin using a 0.005 µm histogram binning interval.

### SPAD calling and cell type comparisons

We ranked smoothed TSA-Seq enrichment scores over all 20 kb bins genome-wide from largest to smallest excluding hg38 unmapped regions, dividing these ranked scores into 100 equal sized groups, defined as percentiles 1 to 100 (lowest to highest scores). Adjacent 20kb bins ranked in the 96-100^th^ percentile were merged to segment these regions as SPADs.

To compare SPADs called in one cell line with TSA-Seq score percentiles in other cell lines, we calculated the mean TSA-Seq score percentile over each SPAD region in each of the other cell lines and plotted these values in a boxplot, with dots showing all values. We used Intervene (Khan and Mathelier 2017) (version 0.6.4) to generate a 4-way Venn diagram using 20-kb binned .bed files of the four cell lines.

### Identification of genomic domains that show different nuclear positions relative to speckles in different cell lines

To identify genomic regions that change nuclear position relative to speckles in cell line pair-wise comparisons, we adapted a previously published method (Peric-Hupkes et al. 2010) that compares the variability of scores in two cell lines with the variability observed in biological replicates. The null hypothesis tested is whether a region is statistically different from the values of the biological replicates of the two different cell lines. The detailed statistical method is provided in **Supplementary methods.**

To identify genomic regions that showed statistically significant changes in position, we set a cutoff p-value of 0.01 and identified all 20kb bins with p-values < 0.01 for either ordering (bigger TSA-Seq values in one cell line or the other). For each ordering, we merged the identified adjacent bins to call domains that changed location. We set a second cutoff to call only domains corresponding to 100 kb or larger.

We defined SPADs as regions with mean TSA-Seq score in the 96-100^th^ genomic percentile. We then identified those SPAD regions with either significantly increased TSA-Seq scores in one cell line within this top 5^th^ percentile range or SPAD regions which showed statistically significant changes between cell lines but which were classified as non-SPAD in one of the two cell lines compared.

### Profiling local TSA-Seq signals and histone marks for changed domains

We used the deepTools suite (Ramirez et al. 2016) (version 3.2.1) to profile TSA-Seq score percentiles and CUT&RUN-Seq log2-fold changes for all the changed regions we identified comparing H1 vs HFFc6. The computeMatrix tool (deepTools) was used to compute signals 2-Mbp upstream and downstream of each region center (10-bp bin). The generated matrix file was used as input for the plotHeatmap tool (deepTools) to plot a heatmap and profile of mean scores for all domains. CUT&RUN-Seq data were generated by the Steven Henikoff laboratory (Seattle, WA) using an automated platform (Janssens et al. 2018). CUT&RUN-Seq data processing is described in **Supplementary methods**.

### RNA-Seq data processing

We applied the ENCODE RNA-Seq processing pipeline (https://github.com/ENCODE-DCC/rna-seq-pipeline) to process RNA-Seq data (Supplementary Table 5). Briefly, we used STAR (Dobin et al. 2013) (version 2.5.1b) to map raw sequencing reads to the human reference genome (https://www.encodeproject.org/files/ENCFF742NER/). Next, we used RSEM (Li and Dewey 2011) (version 1.2.26) to quantify gene expression using the ENCODE gene annotation file (URL: https://www.encodeproject.org/files/ENCFF940AZB/). We used the mRNA fragments per kilobase of transcript per million mapped reads (FPKM) for each gene reported by RSEM for downstream analysis. We generated Reads per million (RPM) tracks for each RNA-Seq dataset using the STAR parameter “--outWigNorm RPM”.

See **Supplementary methods** for correlations of RNA-Seq with TSA-Seq data, differential expression analysis, and gene ontology analysis.

### Estimating nuclear speckle mean distance changes between H1 vs K562 using TSA-Seq

We first correlated the scaled TSA-Seq score in each 20 kb genomic bin in K562 cells with estimated mean nuclear speckle distances (Methods, TSA distance prediction), generating a score-distance dictionary. Using mean scaled TSA-Seq scores for each changed domain in K562 or H1 cells, we then converted these into distances based on this score-distance dictionary for K562 cells. We then calculated the changes in distance predicted in K562 cells for shifts in scaled TSA-Seq scores corresponding to the shift observed comparing scaled scores in H1 versus K562.

### PRO-Seq data processing

We downloaded PRO-Seq data for K562 cells cultured in optimal growth conditions (37 °C, NHS) or treated with heat shock at 42°C for 30 mins (HS30) (GEO: GSE89230), combining raw sequencing reads from two replicates for each condition. We mapped the reads to the human reference genome hg38 (excluded Chromosome Y) using Bowtie2 (Langmead and Salzberg 2012) (version 2.1.0) with default parameters and the rmdup command from SAMtools (Li et al. 2009) (version 1.5) to remove potential PCR duplicates. We then used bamCoverage from the deepTools suite (Ramirez et al. 2016) (version 3.2.1) to normalize the alignment files using BPM as normalization method binning at 50 bp with strand separation: BPM (per bin) = number of reads per bin / sum of all reads per bin (in millions).

### 3D immuno-FISH

FISH probes were made from human BACs (ordered from Invitrogen, RP11-479I13 for *HSPA1A* locus and RP11-173P16 for *HSPH1* locus), as previously described (Chen et al. 2018). K562 cells were plated on poly-L-lysine (Sigma-Aldrich P4707, 70,000-150,000 M.W., 0.01% w/v) coated coverslips (Fisher Sci 12-545-81) with 0.5 – 1 mL at 0.5 – 0.7 million/mL cell density. Cells were incubated in the 37 °C incubator for 10 mins for attachment. IMR90, WI38, Tig3 and HCT116 cells were plated on coverslips (Fisher Sci 12-545-81) and cultured for 24 – 48 hrs to reach a confluence of 60% - 80%. Cells were either moved to water bath for heat shock treatment at 42 °C or kept in the 37 °C incubator as control. 3D immuno-FISH was done as previously described (Chen et al. 2018) with modifications for *HSPA1A* FISH in Fig. S11A-B and *HSPH1* FISH: 1) cells were permeabilized with 0.1% Triton X-100 in high magnesium buffer (5mM MgCl2, 50 mM PIPES (pH 7.0)) instead of PBS before fixation; 2) rabbit anti-SON polyclonal antibody (Pacific Immunology Corp, custom-raised, 1:1000 diluted) was used as the primary antibody.

### Combined, sequential RNA- and DNA- FISH

Since we could not preserve the RNA-FISH signal during the high temperature denaturation in the DNA-FISH procedure, we applied a sequential FISH procedure for the *HSPH1* locus. We conducted single molecule RNA (smRNA)-FISH, recorded images, and then conducted DNA-FISH, found the same cells, and recorded images. Experimental details and methods for measurement and analysis of FISH data are provided in **Supplementary methods.**

### Data access

All TSA-Seq data are available at the 4DN <u>Data Portal</u> (https://data.4dnucleome.org/). See Supplementary Table 9 for a summary and links.

### Code availability

TSA-Seq normalization software is available at https://github.com/zocean/Norma. Codes for all genomic data analyses are available at https://github.com/lgchang27/TSA-Seq-2020.

## Acknowledgements

We thank the UIUC Biotechnology center for guidance with preparation of sequencing libraries and quality control. We thank Drs. K.V. Prasanth, William Brieher, Lisa Stubbs and Huimin Zhao (UIUC, Urbana, IL) for helpful suggestions. We thank Derek Janssens (Henikoff lab, Fred Hutchinson Cancer Research Center, Seattle, WA) for generating and sharing CUT&RUN-Seq data. We thank Belmont lab members for sharing reagents and providing suggestions. We thank members of the 4D-Nucleome Consortium and the Belmont NOFIC U54 Center for helpful suggestions and feedback. This work was supported by National Institutes of Health grant R01 GM58460 (ASB) and U54 DK107965 (ASB, JM).

## Author contributions

LZ designed and performed all experiments, collected data, and conducted microscopy image analysis with ASB’s guidance. LZ and YZ analyzed genomic data with guidance from ASB and JM. YC developed and standardized TSA-Seq 1.0 protocols and provided suggestions for development of TSA-Seq 2.0. OG contributed to the “hybrid” distance-calibration mapping approach. YW processed the raw sequencing data of CUT&RUN-Seq. LZ and ASB wrote the manuscript with critical suggestions from other co-authors. ASB supervised the overall study.

## Notes

https://data.4dnucleome.org/browse/?experimentset_type=replicate&type=ExperimentSetReplicate&award.project=4DN&experiments_in_set.experiment_type.display_title=TSA-seq&experiments_in_set.biosample.biosource.individual.organism.name=human

### Competing Interest Statement

The authors have declared no competing interest.

### Summary of Updates

We have expanded this manuscript to a full-length research article, including more analysis of previous data and also including new heat shock data and analysis. The emphasis is now placed on the biological results- comparing genome organization relative to nuclear speckles in different cell lines and after heat shock- as opposed to the new TSA-Seq 2.0 method per se.

