## Supplementary Methods and Figures for "TSA-Seq reveals a largely “hardwired” genome organization relative to nuclear speckles with small position changes tightly correlated with gene expression changes"

**Supplemental methods**

**Cell culture**

K562 cells were cultured in RPMI-1640 medium supplemented with 10% FBS (Sigma-Aldrich F2442) and 1% 100X Antibiotic-Antimycotic (GIBCO). Cells were seeded at 0.1 million/mL density and passaged or harvested at around 0.75 million/mL density. H1 cells were cultured in Matrigel (Corning 354277, Lot #7128002)- coated flasks in mTeSR medium (STEMCELL Tech 85850). Cells were passaged (or harvested) at optimal density and colony size (see SOP) every 4 or 5 days with 1 in 10 to 1 in 20 splits by digesting colonies into 50-200 μm aggregates using ReLeSR (STEMCELL Tech 05872). HFFc6 cells were cultured in DMEM medium supplemented with 20% heat-inactivated FBS (VWR 97068-091, Lot #035B15). Cells were passaged (or harvested) at 70%-80% confluency without significant drop of mitotic cell ratio with 1 in 2 splits using Trypsin-EDTA (0.05%) (Fisher Sci 25300054). HCT116 cells were cultured in McCoy's 5A Medium supplemented with 10% heat-inactivated FBS (VWR 97068-091, Lot #035B15). Cells were passaged (or harvested) at 70%-80% confluency (around 0.4 million cells/$\mathrm{cm}^2$) and seeded at $4-5 \times 10^4$ cells/$\mathrm{cm}^2$ using Trypsin-EDTA (0.05%) (Fisher Sci 25300054).

**Coverslip TSA staining**

K562 cells were plated on poly-L-lysine (Sigma-Aldrich P4707, 70,000-150,000 M.W., 0.01% w/v) coated coverslips (Fisher Sci 12-545-81) with 0.3-0.5 mL at 0.4-0.7 million/mL cell density. Cells were cultured for 30 mins for attachment. HFFc6 cells were plated on coverslips 1 or 2 days before experiments and harvested at ~80% confluency. Cells were fixed with 1.6% freshly-made paraformaldehyde (PFA) (Sigma-Aldrich P6148) in PBS at room temperature (RT) for 20 mins. Cells were then permeabilized with 0.5% Triton X-100 (Sigma-Aldrich T8787) in PBS (0.5% PBST) at RT for 30 mins, treated with 1.5% $\mathrm{H_2O_2}$ in PBS at RT for 1 hr to quench endogenous peroxidases, and rinsed with 0.1% Triton X-100 (Sigma-Aldrich T8787) in PBS (0.1% PBST) 3x at RT. Cells were blocked with 5% normal goat serum (Sigma-Aldrich G9023) in 0.1% PBST (GS blocking buffer) at RT for 1 hr and then incubated with rabbit anti-SON polyclonal antibody(Chen et al. 2018) (Pacific Immunology Corp, custom-raised) 1:2000 in GS blocking buffer at RT for 5 hrs. Cells were then washed with 0.1% PBST at RT 3 x 5 mins and incubated with HRP conjugated goat anti-rabbit polyclonal antibody (Jackson Immuno 111-035-144) 1:1000 in GS blocking buffer at RT for 5 hrs or at 4 °C for 10-12 hrs. Cells were then washed with 0.1% PBST at RT for 3 x 5 mins and subject to TSA labeling.

For Condition A, the reaction solution is 50% sucrose (w/v), 1/10000 tyramide-biotin or tyramide-FITC (v/v), and 0.0015% $H_2O_2$ (v/v) in PBS. Tyramide-biotin was prepared as previously described(Chen et al. 2018). Tyramide-FITC was prepared according to an online protocol (http://wiki.xenbase.org/xenwiki/index.php/Flourescin_Tyramide_Synthesis). Labeling is done at RT for 10 mins. For Condition E, the reaction solution is 50% sucrose, 1/300 tyramide-biotin and 0.0015% $H_2O_2$ in PBS. Labeling is done at RT for 30 mins. For both conditions, 500 µL reaction solution was applied per coverslip.

For “two-rounds” of TSA labeling, after the 1st-round of TSA, cells were washed with 0.1% PBST at RT for 3 x 5 mins and then subject to a 2nd-round of TSA. After TSA labeling, cells were washed with 0.1% PBST at RT for 3 x 5 mins, stained with Streptavidin-Alexa Fluor 594 (Invitrogen) 1:200 and goat anti-rabbit – Alexa Fluor 647 (Jackson Immuno) 1:200 in GS blocking buffer at RT for 2 hrs or at 4 °C for 10-12 hrs, and then washed with 0.1% PBST at RT for 3 x 5 mins. Coverslips were mounted in DAPI containing, anti-fading medium (0.3 µg/ml DAPI (Sigma-Aldrich)/10% w/v Mowiol 4-88(EMD Millipore)/1% w/v DABCO (Sigma-Aldrich)/25% glycerol/0.1 M Tris, pH 8.5).

**TSA-Seq**

The TSA-Seq procedure was modified from our previous publication (Chen et al. 2018).

***For suspension cells (K562)***, cells were fixed by adding 8% freshly made PFA in PBS to reach a final concentration of 1.6% and incubated at RT for 20 mins. Aldehyde groups were quenched by adding 1.25M (10x) glycine in PBS and mixing at RT for 5mins. Cells were permeabilized with 0.5% PBST at RT for 30 mins, centrifuged at 116 g, and re-suspended in PBS. $H_2O_2$/PBS was added to reach a final $H_2O_2$ 1.5% concentration to quench endogenous peroxidases in a volume of 1 mL per 3 million cells; the cell suspension was incubated by slowly nutating at RT for 1 hr (open tubes 2 or 3 times during the incubation to release the generated gas). Cells were rinsed 3x with 0.1% PBST, blocked with 5% normal goat serum (Sigma-Aldrich G9023) in 0.1% PBST (GS blocking buffer) in a volume of 1 mL / 10 million cells at RT for 1 hr, and then incubated with rabbit anti-SON polyclonal antibody (Chen et al. 2018) (Pacific Immunology Corp, custom-raised) 1:2000 in GS blocking buffer at 1mL / 10 million cells at 4 °C for 20-24 hrs. Cells were then washed with 0.1% PBST at RT for 3 x 5 mins and incubated with HRP conjugated goat anti-rabbit polyclonal antibody (Jackson Immuno 111-035-144) 1:1000 in blocking buffer in a volume of 1 mL / 10 million cells at 4 °C for 20-24 hrs. Cells were washed with 0.1% PBST for 3 x 5 mins, washed with PBS for 5 mins at RT, and then subjected to TSA labeling. Cells were resuspended in 50% sucrose/PBS and then the same volume of 50% sucrose/PBS containing tyramide-biotin and hydrogen peroxide was added to reach the final concentrations for a specific labeling condition (Fig. S1A). The final volume of reaction solution was 1 mL per 10 million. Cells were gently nutated at RT during the TSA labeling time specific to each condition (Fig. S1A). Cells were then washed at RT with 0.1% PBST 3 x 5 mins and with PBS for 5 mins. For each sample, a small portion of cells were attached to a coverslip for anti-biotin and anti-SON staining to visualize the TSA labeling.  Remaining cells were pelleted and either immediately subjected to genomic DNA isolation or stored at -80 °C for later DNA isolation. K562 cells were lysed with high T-E buffer (10 mM Tris and 10 mM EDTA, pH 8.0) containing 0.5% SDS and 0.2 mg/mL Proteinase K (NEB P8107S).  All centrifugations prior to TSA labeling were low-speed at 116 – 130 g for 5 – 10 mins to preserve cell structure.

For the TSA-Seq mapping of K562 cells upon heat shock, cell flasks were incubated in the 37 °C incubator or in 42 °C water bath for heat shock. One replicate was done with TSA-Seq 2.0 for control and heat shock of 30 mins, 1hr, or 2hrs. A second replicate was done with TSA-Seq 1.0 for control and heat shock of 30 mins.  The heat-shocked or control cells were immediately fixed and proceeded for TSA-Seq procedure as described above.

***For attached cells (H1, HFFc6, HCT116)***, cells were grown in tissue culture flasks and fixed by quickly pouring away growth media, adding freshly made 1.6% PFA in PBS, and incubating at RT for 20 mins. Cells were rinsed with PBS, and then washed/permeabilized with 0.5% PBST at RT for 3 x 5 mins. Free aldehyde groups were quenched with 20 mM glycine in PBS at RT for 3 x 5 mins. Cells were then washed with PBS and incubated with 1.5% $H_2O_2$ in PBS at RT for 1 hr. Cells were rinsed 3x with PBS, blocked with 5% normal goat serum (Sigma-Aldrich G9023) in PBS in a volume of 1mL per 25 $cm^2$ flask surface area at RT for 1 hr, and then incubated with rabbit anti-SON polyclonal antibody (Chen et al. 2018) (Pacific Immunology Corp, custom-raised) 1:2000 in 0.1% PBST in a volume of 1 mL per 25 $cm^2$ at 4 °C for 10-12 hrs. Cells were then washed with 0.1% PBST at RT for 3 x 5 mins and incubated with HRP conjugated goat anti-rabbit polyclonal antibody (Jackson Immuno 111-035-144) 1:1000 in 0.1% PBST in a volume of 1 mL per 25 $cm^2$ at 4 °C for 10-12 hrs. Cells were washed at RT with 0.1% PBST for 3 x 5 mins and with PBS for 5 mins. Cells were TSA-labeled with a working solution of 50% sucrose/PBS containing 0.0015% $H_2O_2$ and the condition-specific tyramide-biotin concentration (Fig. S1A) in a volume of 0.8mL per 10 $cm^2$surface area. Cells were incubated at RT for the condition-specific time (Fig. S1A) and then washed at RT with 0.1% PBST 3 x 5 mins and with PBS for 5 mins. For each staining, a small portion of attached cells were scraped off and loaded onto coverslips for anti-biotin and anti-SON immunostaining to visualize the TSA-labeling. Remaining cells were washed with high T-E buffer (10 mM Tris and 10 mM EDTA, pH 8.0) at RT for 5 mins and lysed with high T-E buffer containing 1% SDS and 0.2 mg/mL Proteinase K. Cell lysates were collected and immediately subjected to genomic DNA extraction. All incubation and washing steps before cell-lysing were done by gently shaking the original flasks.

Cells on coverslips were stained with Streptavidin-Alexa Fluor 594 (Invitrogen) 1:200 and goat anti-rabbit – FITC (Jackson Immuno) 1:500 in GS blocking buffer at RT for 2 hrs or at 4 °C for 10-12 hrs. Cells were then washed with 0.1% PBST at RT for 3 x 5 mins. Coverslips were mounted in DAPI containing, anti-fading medium (0.3 µg/ml DAPI [Sigma-Aldrich]/10% w/v Mowiol 4-88 [EMD Millipore]/1% w/v DABCO [Sigma-Aldrich]/25% glycerol/0.1 M Tris, pH 8.5).

Genomic DNA was extracted by phenol/chloroform as previously described (Chen et al. 2018). For samples with low DNA concentration, glycogen (Roche) was added to a final concentration of 0.05 mg/mL to facilitate ethanol precipitation. Isolated DNA was fragmented to 100-600 bp using a Bioruptor Pico (Diagenode) machine with a mode of 30 sec ON – 30 sec OFF. Inspection of DNA gels determined the number of required sonication cycles. Biotin labelled DNA fragments were isolated with streptavidin beads as previously described (Chen et al. 2018).

Sequencing libraries were constructed using the TruSeq ChIP Sample Prep Kit (Illumina, IP-202-1012) for H1, HCT116, HFFc6 samples, or the Hyper Library Construction Kit (Kapa Bio) for K562 Condition A, B, C samples, or a protocol developed in the John Lis laboratory (Cornell

University) for K562 Condition D, E samples. The John Lis lab protocol includes steps of end repair (End-It DNA End-Repair Kit: Epicentre Biotech ER0720), 3’-end A tailing (Klenow Fragment: NEB M0212S), and indexed adaptor ligation (NEXTflex ChIP-Seq Barcodes-6: Bioo Sci 514120; T4 DNA Ligase [Rapid]: Enzymatics Inc L6030-HC-F). Libraries were amplified by 8-12 PCR cycles. qPCR was used to measure the concentrations of each library which were then pooled at equimolar concentration for each lane and sequenced for 101 cycles from one end of the fragments on a HiSeq4000 using a HiSeq4000 sequencing kit version 1 (Illumina). Fastq files were generated and demultiplexed with the bcl2fastq Conversion Software (Illumina).

**Dot blot and DNA biotinylation estimation**

Biotinylation levels of DNA were assayed by dot blot prior to biotin-avidin pulldown of DNA. 5-fold serial dilutions of a biotin end-labeled 250 bp fragment were used as biotin concentration standards. The PCR product was produced from *Drophophila* DNA cloned within a BAC (BACR48E12) using primers: GAAACATCGC/iBiodT/GCCCATAAT (forward) and AGAAGCAGCTACGCTCCTCA (reverse) resulting in 1 biotin per fragment. These PCR standards were combined with control, unbiotinylated, sonicated K562 genomic DNA (100-600 bp) to reach a final DNA concentration the same as the test DNA sample (400 or 600 ng/µL) to eliminate concentration effects on DNA crosslinking to membranes.

DNA was spotted with 1 or 1.5 µL (same volume for all standards and samples in a specific experiment) onto a nitrocellulose membrane (0.45 µm; Bio-Rad), and were UV cross-linked to the membrane (0.24J; UV Stratalinker 2400; Agilent Technologies). The membrane was blocked with SuperBlock (TBS) blocking buffer (Thermo) with 0.05% Tween-20 (Fisher) at RT for 1 hr, and then incubated with streptavidin-HRP (Invitrogen 43-4323) diluted 1:10000 in blocking buffer at 4 °C for 1-3 hr or overnight. The membrane was then washed with 0.05% Tween-20 in

TBS at RT for 6 x 5 mins with rigorous shacking. The membrane was treated with SuperSignal West Femto chemiluminescent substrate (Thermo) and developed with HyBlot CL film (Denville) or in an iBright machine (Invitrogen). Sample biotinylation level (kilobases of DNA with one biotin labeling on average) was calculated following the equation:

$$\frac{1}{l_{std}} \times c_{std} = \frac{1}{l_{smp}} \times c_{smp}$$

$l_{std}$: standard DNA length per biotin (0.25 kb). $l_{smp}$: average sample DNA length (kb) per biotin (to be calculated). $c_{std}$: concentration of the 0.25kb standard DNA that has the same signal intensity with the tested sample DNA. $c_{smp}$: concentration of the tested sample DNA.

**TSA-Seq normalization**

TSA-Seq normalization followed a similar scheme as used in the previous paper (Chen et al. 2018), using matched pulldown and input data with the following updates: First, instead of using sliding windows, we used separate genomic bins of 20kb. Each mapped read is exclusively assigned to a window according to its largest aligned position in the reference genome. Second, we modified the calculation of TSA-Seq enrichment score, as described in the next paragraph.

$N_{TSA}$ and $N_{input}$ represent the original number of mapped reads in each 20kb bin in the pulldown and input samples, respectively. $N_{TSA}$ in each bin (bins without mapped reads are skipped) was normalized by input read number, according to equation (1), where $Ave(N_{input})$ is the average value of $N_{input}$ across genome-wide windows calculated by dividing the genome-wide sum of $N_{input}$ by the number of bins containing mapped reads:

$$N'_{TSA} = \frac{N_{TSA} \times Ave(N_{input})}{N_{input}} \qquad (1)$$

The TSA-Seq enrichment score is then defined as the log2 ratio between $N'_{TSA}$ and the genome-wide average of $N'_{TSA}$ ($Ave(N'_{TSA})$, calculated using the number of bins containing non-zero numbers of mapped reads, as per equation 2:

$$\text{TSA-Seq enrichment score} = \log_2\left(\frac{N'_{TSA}}{Ave(N'_{TSA})}\right) \quad (2)$$

This normalized TSA-Seq enrichment score can be considered as the log2 ratio of relative enrichment or depletion of DNA in a specific bin relative to the average pulldown value.

For subsequent analyses, the non-overlapping 20kb binned signals were smoothed by convolution using a Hanning window of length 21 (21 × 20 kb or 420 kb).

K562 pulldown data from all five conditions were normalized using one input constructed from fragmented K562 genomic DNA. H1, HCT116 and HFFc6 data were normalized using separate input libraries made from the DNA used for pulldown for each TSA-labeling experiment.

**“Hybrid” method for TSA distance prediction**

To get the calibration equation $y = y_0 + Ae^{R_0 x}$, first we obtained the exponential decay parameter $R_0$ by fitting 16 FISH measurements from our previously published data (Chen et al. 2018) to the new TSA-Seq data (Supplementary Table 1). For each FISH probe, we used the previously published mean cytological distance to speckles based on measurements of 100 alleles (Chen et al. 2018) (16 probes, Supplementary Table 1). We used the smoothed TSA-Seq data and calculated the mean TSA-Seq enrichment values over the genomic regions cloned within the BACs used to generate FISH probes. We fit the 16 TSA-Seq fold-enrichment values, y, and their corresponding mean speckle distances, x, to the exponential function $y = y_0 + Ae^{R_0 x}$ using OriginPro software (OriginLab) to obtain the exponential parameter, $R_0$.

Next, we obtained $y_0$ and $A$ based on the minimum and maximum TSA-Seq fold-enrichment values, $y_{min}$ and $y_{max}$: $\lim_{x \to \infty (x_{\max})} (y_0 + Ae^{R_0 x}) = y_0 = y_{min}$ $(R_0 < 0)$ and $\lim_{x \to 0} (y_0 + Ae^{R_0 x}) = y_0 + A = y_{max}$. With the parameters $R_0$, $y_0$ and $A$ (Supplementary Table 2), we converted TSA-Seq enrichment values from all 20 kb bins of smoothed TSA-Seq data into speckle distances by computing the inverse equation for x in the exponential function: $x = \frac{1}{R_0} \ln \frac{y - y_0}{A}$.

**Statistical method to identify changed genomic bins in pair-wise cell type comparison**

We adapted a previously published method (Peric-Hupkes et al. 2010) to test whether a genomic bin (20kb) is statistically different from the values of the biological replicates of the two different cell lines.

For each dataset, we first rescaled TSA-Seq enrichment scores (20kb bin) linearly between their min and max values to a new 1-100 scale based on equation (3) and rounded up to integers with min assigned as 1 instead of 0.

$$\text{Scaled enrichment score (bin i)} = \frac{\text{TSA-Seq enrichment score (bin i)} - \text{min}}{\text{max} - \text{min}} \times 100 \qquad (3)$$

To reduce the influence of outliers on this rescaling, we used a large and a small percentile of all ranked values as the max and min values, respectively (e.g. 99.95$^{th}$ and 0.05$^{th}$ percentile for HFFc6 and H1 comparison).

For a pair-wise comparison between two cell lines, we rescaled the SON TSA-Seq scores for two biological replicates for each cell line:

$$\text{TSA}_{\text{H1}_{rep1}}, \text{TSA}_{\text{H1}_{rep2}}, \text{TSA}_{\text{HFFc6}_{rep1}}, \text{TSA}_{\text{HFFc6}_{rep2}}$$

For example, $\mathrm{TSA}_{\mathrm{H1}_{rep1}}$ denotes the H1 TSA-Seq biological replicate 1 and is a row vector of N values from all 20kb non-overlapping bins with mapped reads in the genome. Scatterplots show near uniform data noise across the genome for both cell lines.

We averaged the replicates for the same cell line and used the residual, Δ, between two cell lines for comparison:

$$\Delta = \frac{1}{2}\{ ( \mathrm{TSA}_{\mathrm{HFFc6}_{rep1}} + \mathrm{TSA}_{\mathrm{HFFc6}_{rep2}} ) - ( \mathrm{TSA}_{\mathrm{H1}_{rep1}} + \mathrm{TSA}_{\mathrm{H1}_{rep2}} ) \} \quad (4)$$

We define data variance as the difference between biological replicates for the same cell line. We averaged the variance between the two cell lines to be compared to construct a vector ϕ including all possible orderings with a length of 4N, where N= number of genomic bins excluding unmapped regions from hg38 genome:

$$\Phi = \frac{1}{2}\{ ( \mathrm{TSA}_{\mathrm{H1}_{rep1}} - \mathrm{TSA}_{\mathrm{H1}_{rep2}} ) + ( \mathrm{TSA}_{\mathrm{HFFc6}_{rep1}} - \mathrm{TSA}_{\mathrm{HFFc6}_{rep2}} ),$$

$$( \mathrm{TSA}_{\mathrm{H1}_{rep1}} - \mathrm{TSA}_{\mathrm{H1}_{rep2}} ) + ( \mathrm{TSA}_{\mathrm{HFFc6}_{rep2}} - \mathrm{TSA}_{\mathrm{HFFc6}_{rep1}} ),$$

$$( \mathrm{TSA}_{\mathrm{H1}_{rep2}} - \mathrm{TSA}_{\mathrm{H1}_{rep1}} ) + ( \mathrm{TSA}_{\mathrm{HFFc6}_{rep1}} - \mathrm{TSA}_{\mathrm{HFFc6}_{rep2}} ),$$

$$( \mathrm{TSA}_{\mathrm{H1}_{rep2}} - \mathrm{TSA}_{\mathrm{H1}_{rep1}} ) + ( \mathrm{TSA}_{\mathrm{HFFc6}_{rep2}} - \mathrm{TSA}_{\mathrm{HFFc6}_{rep1}} ) \} \quad (5)$$

The vector Φ serves as the null distribution against which to test single bin value difference from Δ for statistical significance. Φ can be viewed as a set of observations with random variable ϕ, which is approximately Gaussian distributed with parameters μ and $\sigma^2$.

We test if a single element in Δ (a 20kb bin score) shows a significant change between the two cell lines to be compared by comparing with ϕ. We calculated p-values for both tails of the distribution separately: for equation (4), we separate the bins with values above or below 0 to obtain bins with bigger TSA scores in either HFFc6 or H1. We calculated p-values for all 20kb bins and displayed -log10 (p-value) in the genomic tracks.

**CUT&RUN-Seq data processing**

CUT&RUN-Seq data in H1 and HFF were downloaded from the 4DN data portal (https://data.4dnucleome.org/); these were generated by the Steven Henikoff laboratory (Seattle, WA) using an automated CUT&RUN platform (Janssens et al. 2018). First, we trimmed the paired end reads with trimmomatic (Bolger et al. 2014). The parameters used were ILLUMINACLIP:$adapter/TruSeq3-PE-2.fa:2:15:4:4:true LEADING:20 TRAILING:20 SLIDINGWINDOW:4:15 MINLEN:25. Then we used bowtie2 (Langmead and Salzberg 2012) to align the trimmed reads to reference genome hg38. We set the minimum fragment length to be 10 and maximum fragment length to be 700. Then we sorted and removed duplicate reads with SAMtools (Li et al. 2009). Finally, we used MACS2 (Zhang et al. 2008) to generate fold enrichment (FE) tracks and perform peak calling.

**Correlating RNA-Seq with TSA-Seq data**

To correlate TSA-Seq score with gene expression, we ranked TSA-Seq enrichment scores of genome-wide 20kb bins (bins without mapped reads were removed) from largest to smallest, divided them into 20 equal sized groups using the cut function in R, and named them as vigintiles from vigintile 1 to vigintile 20 (smallest to largest). For each protein-coding gene (based on the GENCODE annotation version 24), we calculated the average TSA-Seq enrichment score across the whole gene region and assigned the gene to the corresponding TSA-Seq vigintile group according to the TSA-Seq enrichment score ranges for each vigintile. RNA-Seq analysis and TSA-Seq correlation results were summarized in Supplementary Table 6.

A housekeeping gene list was downloaded from https://www.tau.ac.il/~elieis/HKG/HK_genes.txt (Eisenberg and Levanon 2013) and processed as previously described (Chen et al. 2018) but using hg38 RefSeq gene annotation. 3791 protein-coding genes were identified as housekeeping genes, and the remaining 16541 protein-coding genes were determined as non-housekeeping genes.

To correlate repositioned regions with expression differences between two cell lines, first we identified all genes located within these genomic regions by overlapping gene and region coordinates (the whole gene must be located within the region to be called). Then we calculated the log2 ratio between the gene FPKM values in the two cell lines and plotted these log2 ratios against the region mean scaled TSA-Seq score (max-min normalized, 1-100) change between the two cell lines.

For differential expression analysis between H1 and HFF (RNA-Seq datasets summarized in Supplementary Table 5), we used STAR (Dobin et al. 2013) (version 2.5.3a) to map the raw sequencing reads using the index files from the ENCODE project (URL: https://www.encodeproject.org/files/ENCFF742NER/). Next, we used htseq-count function of HTSeq (Anders et al. 2014) (version 0.9.1) to count raw read number for each gene with the gencode.v24 annotation (https://www.encodeproject.org/files/gencode.v24.primary_assembly.annotation/). To identify differentially expressed (DE) protein-coding genes between H1 and HFF, we used DEseq2 (Love et al. 2014) (version 1.24.0) with thresholds for adjusted P-value of <0.01 and for fold-expression change of >2-fold (Results summarized in Supplementary Table 7). For DE genes with significantly higher expression in one cell line versus the other, we identified which of these genes located entirely within "repositioned" domains that are located closer to nuclear speckles in this cell line versus the other by overlapping gene genomic position with the coordinates of the domains that showed significantly higher scaled TSA-Seq scores in this cell line. The remaining DE genes with significantly higher expression in this cell line were determined as differentially expressed but not located within domains that reposition closer to speckles ("non-repositioned").

Next we calculated the mean scaled TSA-Seq scores (max-min normalized, 1-100) across each of the genes and generated scatter plots to show the correlation of these TSA-Seq scores between the two cell lines. We calculated log2-fold expression changes for repositioned versus non-repositioned DE genes, and plotted boxplots of these expression changes for all genes in each of these two categories. We used DAVID (Huang da et al. 2009b; Huang da et al. 2009a) (version 6.8) to conduct gene ontology (GO) analysis using the "GOTERM_BP_DIRECT" category (Results summarized in Supplementary Table 8). We compared GO terms for the repositioned and non-repositioned DE genes by plotting bar plots of -log10 P-values of top 5 terms sorted by P-values.

**Combined (sequential) RNA- and DNA- FISH**

We first conducted smRNA-FISH, acquired images, and then conducted DNA-FISH. To return to the same cells to record DNA-FISH images, we recorded cell coordinates relative to etched grids by using gridded glasses from Ibidi µ-Dishes (35 mm, high Glass Bottom, Grid-50; Ibidi 81158) OR relative to etched reference points on coverslips. For the latter approach, we etched three reference points on a regular glass coverslip (Fisher Sci 12-545-81) with a diamond pencil. We used the point-marking function in the DeltaVision OMX microscope to record coordinates of these reference points and then the fields of view containing the RNA-FISH images of cells. We then stripped off coverslips from slides, conducted DNA-FISH, and remounted the coverslips. In the second-round of microscopy for DNA-FISH, we first measured locations of the original three reference points. We then established a transformation matrix to transform the coordinates of our cell point list from the first microscopy round into the coordinate system of the reoriented coverslip in the second microscopy round. We then used these new coordinates to reposition the microscope stage to acquire images for the same cells imaged in the first round of microscopy. For the detailed protocol, R code, and example coordinate lists, see https://github.com/lgchang27/Find_cells).

The glasses/coverslips were autoclaved and coated with poly-L-lysine (Sigma-Aldrich P4707, 70,000-150,000 M.W., 0.01% w/v). K562 cells were plated on the coated glasses with 1 mL at 0.4 – 0.7 million/mL cell density and incubated in 37 °C incubator for 10 mins for attachment. Cells were then either moved to water bath for heat shock treatment at 42 °C for 1 hr or kept in the 37 °C incubator as control.

smRNA-FISH probes targeting the 5’ UTR (398 bp) and the first intron (2612 bp) of *HSPH1* gene (Supplementary Table 4) were designed, obtained and labeled by Cy5 fluorophore as previously described (Kim et al. 2019). The smRNA-FISH procedure was modified from that previously described (Kim et al. 2019). Cells were fixed with 4% freshly-made paraformaldehyde (PFA) in PBS at room temperature (RT) for 12 mins. Cells were washed with PBS for 3 x 5 mins and then permeabilized with 0.5% Triton X-100 in PBS (0.5% PBST) containing 2 mM ribonucleoside vanadyl complex (NEB S1402) at RT for 10 mins. Cells were rinsed with PBS for 3 times and then equilibrated with freshly made wash buffer (10% formamide [Sigma-Aldrich F9037] and 2×SSC) at RT for 30 mins. Cells were then incubated with smRNA-FISH probes (final concentration: ~300 – 500 nM) diluted in hybridization buffer (2× SSC, 10% formamide, 10% wt/vol dextran sulfate [Sigma-Aldrich D8906], 1 mg/ml Escherichia coli tRNA [Sigma-Aldrich

R8759], 2 mM ribonucleoside vanadyl complex [NEB S1402], and 0.2mg/mL RNase-free BSA [Ambion AM2618, 50 mg/mL] in water) at 37°C for 15 – 17 hrs. Cells were then washed with wash buffer at 37°C for 2 x 30 min. Cells were then post-fixed with 4% freshly-made paraformaldehyde (PFA) in PBS at room temperature (RT) for 12 mins and washed with PBS for 3 x 5 mins. Cells were blocked with 0.5 mg/mL RNase-free BSA in 0.5% PBST containing 2 mM ribonucleoside vanadyl complex at RT for 30 mins. Cells were incubated with mouse anti-Lamin B1/B2 monoclonal antibody [clone 2D8] (Chen et al. 2018) 1:1000 diluted in PBS containing 0.5 mg/mL RNase-free BSA and 2 mM ribonucleoside vanadyl complex at RT for 2hrs. Cells were washed with PBS for 3 x 5 mins and then incubated with goat anti-mouse – AMCA (Jackson Immuno 115-155-003) 1:50 diluted in PBS containing 0.5 mg/mL RNase-free BSA and 2 mM ribonucleoside vanadyl complex at 4 °C for 8 – 10 hrs. Cells were washed with PBS for 3 x 5 mins. Glasses/coverslips were mounted on slides with a non-solidifying glycerol-based medium (90% glycerol, 0.1xSSC, 0.1% p-phenylenediamine [pH 9]) and sealed with nail polish. All water, PBS and SSC buffer used in the smRNA-FISH and immunostaining procedures described above were pre-treated with diethyl pyrocarbonate (DEPC) to prevent RNase contamination. Images for smRNA-FISH were acquired using a DeltaVision OMX microscope system (GE Healthcare).

After acquiring images, the glasses/coverslips were scratched off carefully from slides and the mounting medium was washed off in 4xSSC/0.1% Triton X-100 42 °C for 3 x 5 mins. Cells were permeabilized with 0.5% PBST at RT for 10 mins, blocked with 5% normal goat serum (Sigma-Aldrich G9023) in 0.1% PBST (GS blocking buffer) at RT for 1 hr, and then incubated with rabbit anti-SON polyclonal antibody (Chen et al. 2018) (Pacific Immunology Corp, custom-raised) 1:1000 and mouse anti-Lamin B1/B2 monoclonal antibody [clone 2D8] (Chen et al. 2018) 1:1000 diluted in GS blocking buffer at 4 °C for 10 hrs. Cells were then washed with 0.1% PBST at RT for 3 x 5 mins and incubated with goat anti-rabbit – Texa Red (Jackson Immuno 111-075-003) 1:250

(or goat anti-rabbit – FITC [Jackson Immuno 111-095-144] 1:200) and goat anti-mouse – AMCA (Jackson Immuno 115-155-003) 1:50 diluted in GS blocking buffer at RT for 5 hrs. Cells were then washed with 0.1% PBST at RT for 3 x 5 mins and continued with 3D DNA-FISH for the *HSPH1* locus as described in main Methods. Glasses/coverslips were finally mounted on slides. The same cells were found as described above to acquire images for immunostaining and DNA-FISH.

**FISH image analysis**

To measure the distance between DNA FISH dot signal and speckle, we took the shortest 3D distance from the center of each FISH signal to the edge (defined as 50% drop from highest speckle intensity to nuclear background intensity) of its nearest nuclear speckle using FIJI. For the elongated DNA FISH signal, we consider the signal as a composition of multiple diffraction-limited dots that are connected. We took the shortest distance between these dots (center) and a nearby speckle (edge) as the signal's distance to speckle. To measure the elongated DNA FISH signal length, we measured the length of the trajectory connecting the dot centers. All DNA-FISH distance and signal length measurements were done with deconvolved images. All OMX images used for distance or signal length measurement were aligned for channel correction by bead calibration using the SoftWoRx software.

To measure the smRNA-FISH intensity, we projected optical sections (sum) that contain the FISH signal in z with the "z projection" function in FIJI. We measured the sum of intensity of the FISH signal area in the z-projected image as the signal intensity. We also measured 5 surrounding areas and took the mean as background intensity. We subtracted the background intensity from the signal intensity to have the normalized RNA-FISH signal intensity. All RNA-FISH images from the same experiment were using the same exposure time and % transmitted exciting light. All RNA-FISH intensity measurements were done with non-deconvolved raw images.

Plotting and statistical analysis for distance, length, and RNA signal intensity were done with R. All figure panel images were prepared using FIJI and Illustrator CC (Adobe). RNA-FISH images were shown with projection of 20 slices (sum) that contain the FISH signal in the middle of z (non-deconvolved). All other images were shown with one slice from 3D image stacks (deconvolved).

http://wiki.xenbase.org/xenwiki/index.php/Flourescin_Tyramide_Synthesis.

Huang da W, Sherman BT, Lempicki RA. 2009a. Bioinformatics enrichment tools: paths toward the comprehensive functional analysis of large gene lists. *Nucleic acids research* **37**: 1-13.

Huang da W, Sherman BT, Lempicki RA. 2009b. Systematic and integrative analysis of large gene lists using DAVID bioinformatics resources. *Nature protocols* **4**: 44-57.

Janssens DH, Wu SJ, Sarthy JF, Meers MP, Myers CH, Olson JM, Ahmad K, Henikoff S. 2018. Automated in situ chromatin profiling efficiently resolves cell types and gene regulatory programs. *Epigenetics Chromatin* **11**: 74.

Kim J, Venkata NC, Hernandez Gonzalez GA, Khanna N, Belmont AS. 2019. Gene expression amplification by nuclear speckle association. *The Journal of cell biology* doi:10.1083/jcb.201904046: jcb.201904046.

Langmead B, Salzberg SL. 2012. Fast gapped-read alignment with Bowtie 2. *Nature methods* **9**: 357-359.

Li H, Handsaker B, Wysoker A, Fennell T, Ruan J, Homer N, Marth G, Abecasis G, Durbin R, Genome Project Data Processing S. 2009. The Sequence Alignment/Map format and SAMtools. *Bioinformatics* **25**: 2078-2079.

Love MI, Huber W, Anders S. 2014. Moderated estimation of fold change and dispersion for RNA-seq data with DESeq2. *Genome biology* **15**: 550.

Peric-Hupkes D, Meuleman W, Pagie L, Bruggeman SW, Solovei I, Brugman W, Graf S, Flicek P, Kerkhoven RM, van Lohuizen M et al. 2010. Molecular maps of the reorganization of genome-nuclear lamina interactions during differentiation. *Molecular cell* **38**: 603-613.

Zhang Y, Liu T, Meyer CA, Eeckhoute J, Johnson DS, Bernstein BE, Nusbaum C, Myers RM, Brown M, Li W et al. 2008. Model-based Analysis of ChIP-Seq (MACS). *Genome biology* **9**: R137.

**Supplementary figures**

**A** **Condition:**

A → (3.3X tyramide conc.) → B → (3X reaction time) → C → (3X tyramide conc.) → D → (3.3X tyramide conc.) → E

| Condition | Sucrose | Tyramide-Biotin | Hydrogen peroxide | Reaction time |
|---|---|---|---|---|
| A | 50% | 1:10000 | 0.0015% | 10 min |
| B | 50% | 1:3000 | 0.0015% | 10 min |
| C | 50% | 1:3000 | 0.0015% | 30 min |
| D | 50% | 1:1000 | 0.0015% | 30 min |
| E | 50% | 1:300 | 0.0015% | 30 min |

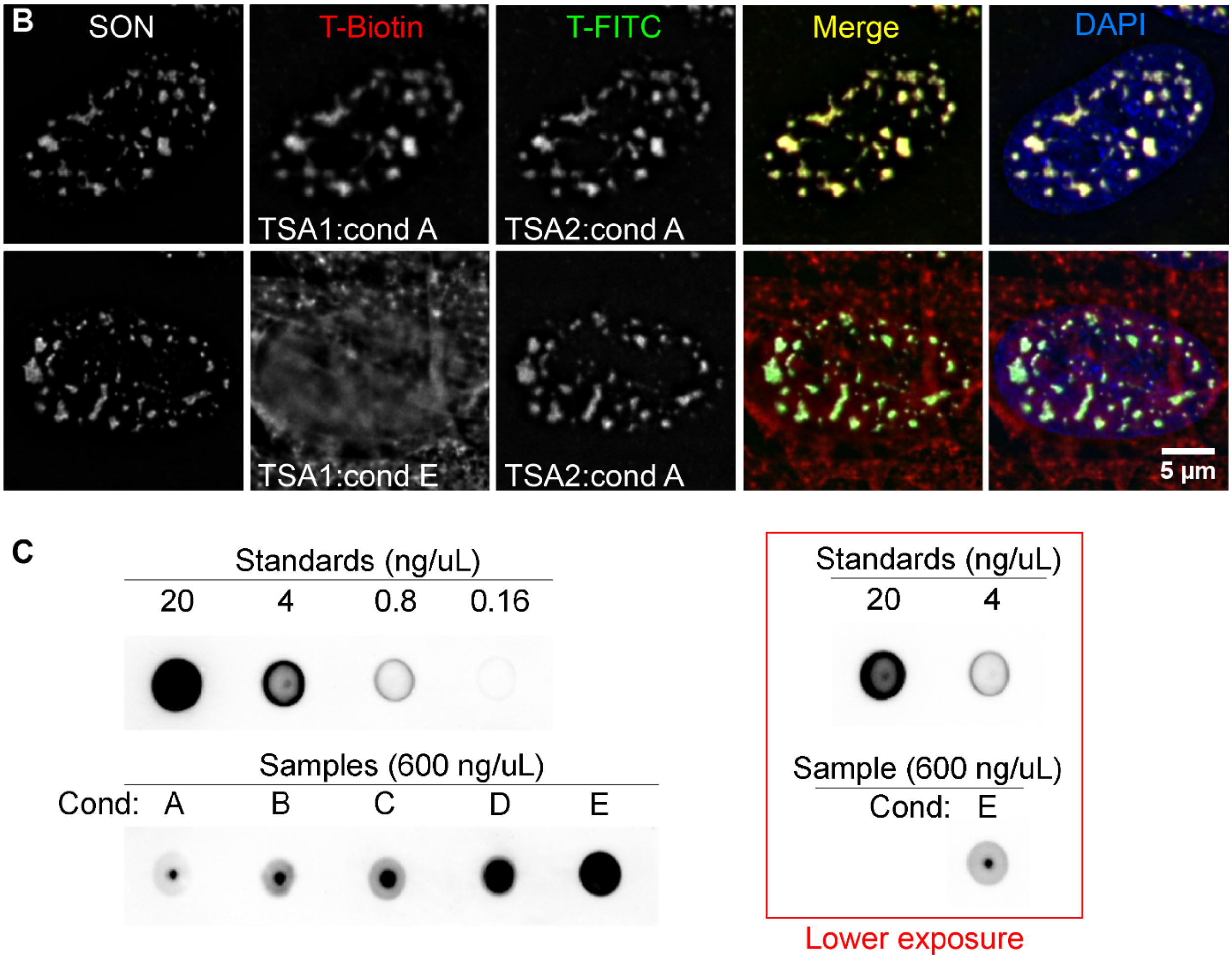

**Supplementary Figure 1. TSA-Seq 2.0 development and evaluation. A)** Experimental design of enhanced TSA labeling conditions. All labeling conditions include 50% sucrose (w/v) and 0.0015% hydrogen peroxide (v/v). From Conditions A to E, serial increases in tyramide-biotin concentration and/or longer reaction times are applied. **B)** Two sequential rounds of SON TSA-labeling demonstrate speckle-specific DNA TSA-labeling after previous "super-saturation" TSA staining in HFFc6. Top row: Two sequential Condition A (non-saturating) rounds of SON TSA-labeling reveals speckle-specific TSA-labeling for both round 1 (tyramide-biotin) and round 2 (tyramide-FITC). Bottom row: First round of SON TSA-labeling using super-saturation Condition E (tyramide-biotin) produces non-specific staining due to saturation of protein-labeling, but second round with tyramide-FITC shows speckle-specific TSA-labeling consistent with sub-saturation DNA labeling: Left to right- SON immunostaining (grey), tyramide-biotin labeling (red), tyramide-FITC (green), merged channels, merged channels plus DAPI-staining (blue). **C)** Dot blot estimation of DNA-biotinylation levels after SON TSA labeling using Conditions A-E. Top: Biotinylated-DNA standards: dilutions of biotinylated PCR 250-bp product (0.16 – 20 ng/µl) containing 1 biotin / DNA molecule combined with unlabeled K562 genomic DNA (100-600 bp fragments) to a total DNA concentration of 600 ng/µl. Bottom: Sonicated genomic DNA (100-600 bp) from K562 cells after SON TSA-labeling using Conditions A-E (left to right). Lower exposure images for first two standards and condition E shown in box.

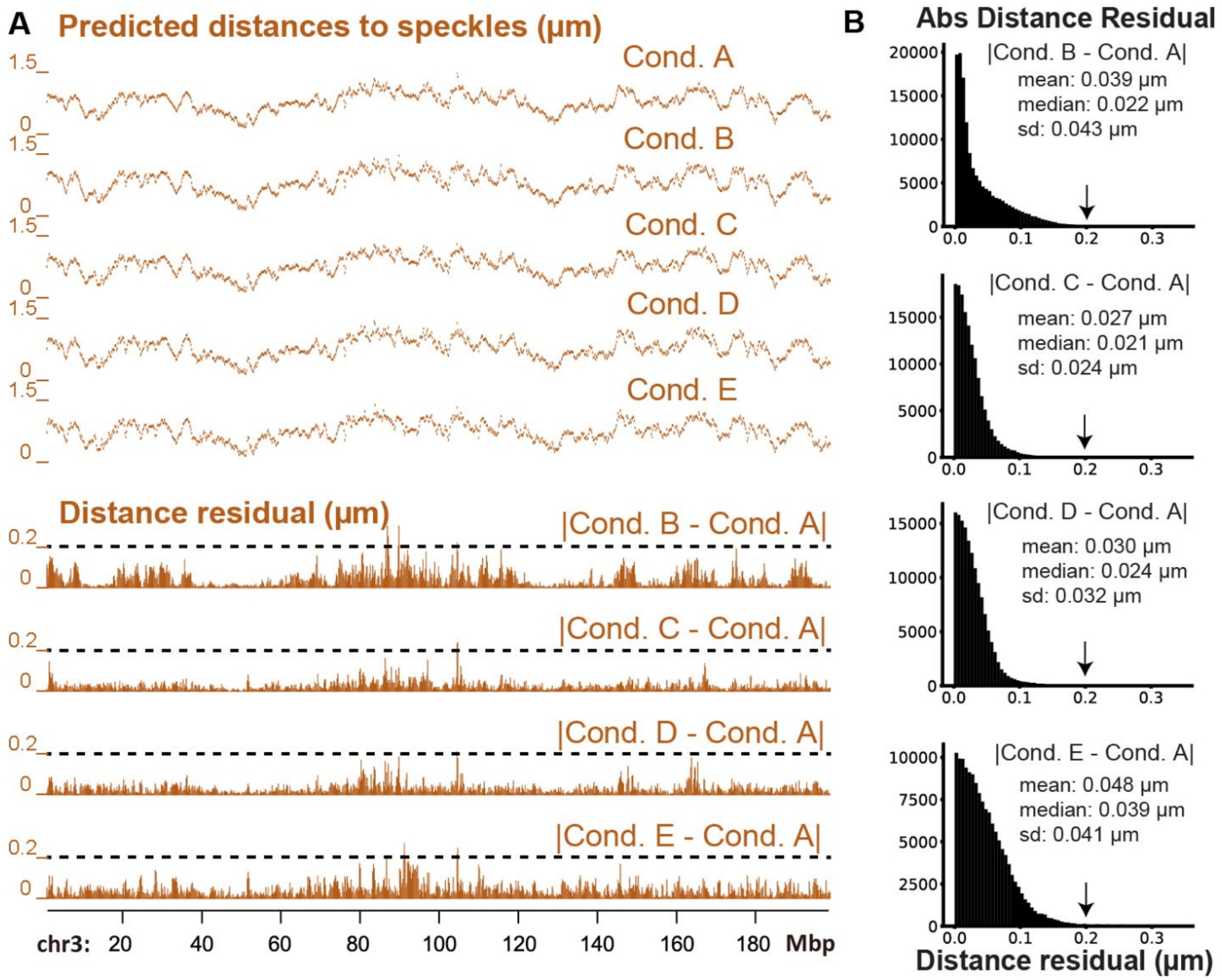


**Supplementary Figure 2. SON TSA Conditions A-E produce quantitatively similar genome-wide estimates of mean distances to speckles. A)** Chromosome 3 tracks showing: (Top) Mean distances (µm) to nuclear speckles in K562 cells estimated from SON TSA-Seq produced using Conditions A-E. (Bottom) Absolute values of distance residuals (µm) between Condition A versus Conditions B-E. **B)** Histograms of absolute value distance residuals between Condition A versus Conditions B-E from all 20 kb bins across the genome. Number (y-axis), distance residual values (x-axis: intervals of 0.005 µm). Arrows mark 0.2 µm value shown as dotted horizontal lines in **A)**.

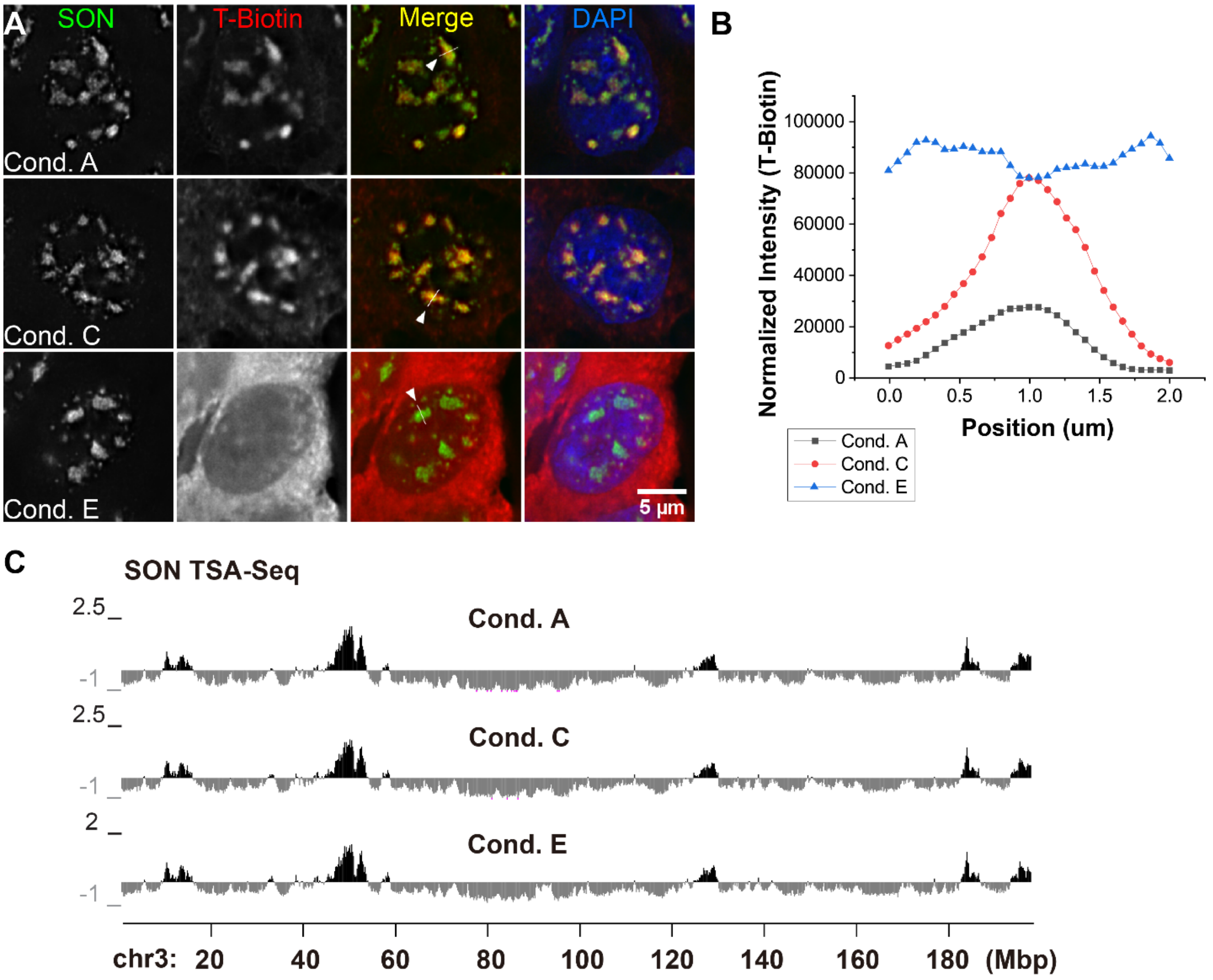


**D**

| Cond. | Cell N (millions) | Pulldown DNA (ng) |
|---|---|---|
| A | 440 | 4.8 |
| C | 220 | 9.3 |
| E | 220 | 59.3 |

**Supplementary Figure 3. SON TSA-Seq using Conditions A-E in HCT116 cells parallels results from K562 cells. A)** Similar changes in reduced specificity of cellular anti-biotin staining after SON TSA in HCT116 cells with increased tyramide labeling using Conditions A (top) versus C (middle) and E (bottom). Left to right: SON immunostaining, streptavidin staining of tyramide-biotin, merged channels (SON, green; biotin, red), plus DAPI (blue). **B)** Tyramide-biotin intensities along line profiles spanning nuclear speckles in **A)** for Conditions A, C, and E. **C)** Similar SON TSA-Seq enrichment score profiles (chromosome 3, 20 kb bins) for Conditions A, C, and E. **D)** Cell numbers used and pulldown DNA yields for Conditions A, C, and E.

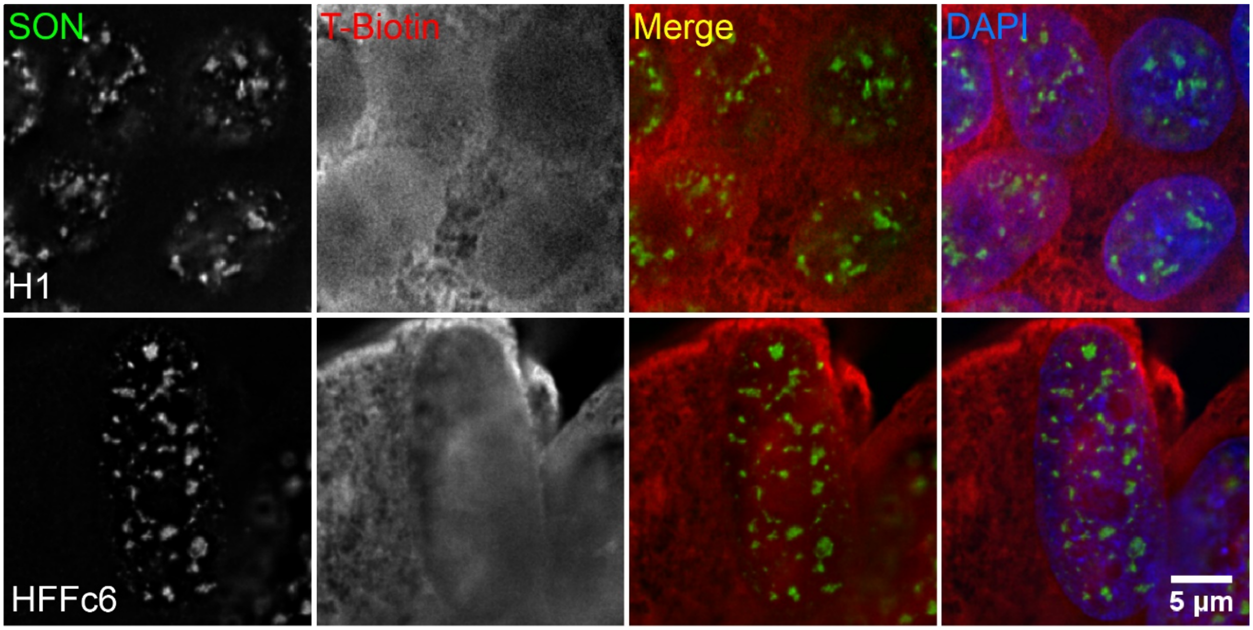


**Supplementary Figure 4. Microscopy assays of biotin-labeling after Condition E SON TSA-labeling in H1 and HFFc6.** Cells were immunostained to verify biotin-labeling after Condition E SON TSA-Seq: (Left to right) SON immunostaining, streptavidin staining of tyramide-biotin, merged image (SON, green; biotin, red), merged image plus DAPI (blue) in H1 (top) and HFFc6 (bottom) cells.

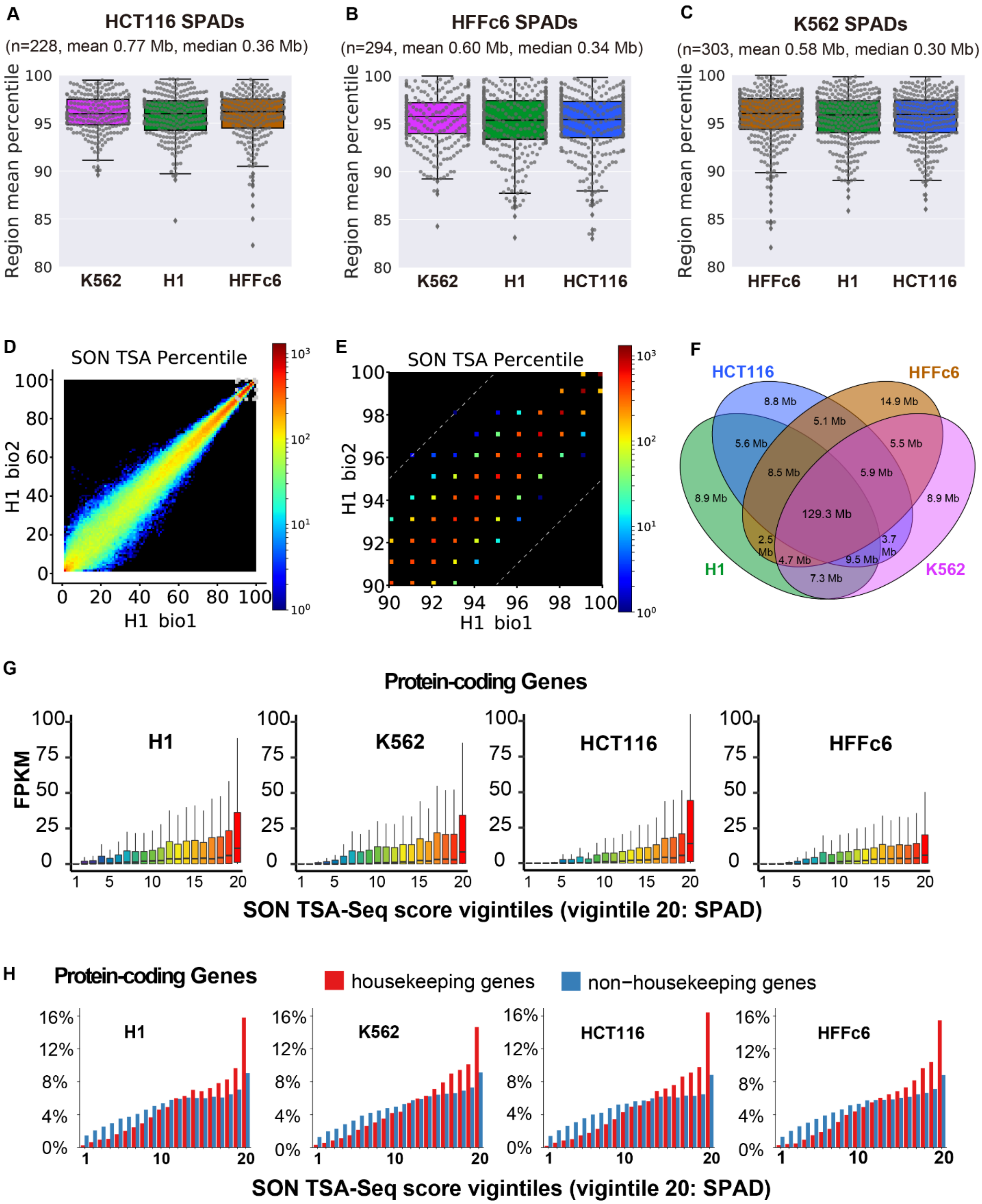


**Supplementary Figure 5. SPADs from one cell line remain high TSA-Seq score percentiles in other cell lines, correlated with high gene expression levels and enriched in house-keeping genes in all cell lines**

**examined. A-C)** Comparison of SPADs from HCT116, HFFc6, and K562 cells with SPADs in other cell lines. TSA-Seq score percentile distributions of SPADs from HCT116 (**A**), HFFc6 (**B**), and K562 (**C**) in the other 3 cell lines. Box plot displays are as described in Fig. 2B legend. **D-E)** 2D histograms show correlation of SON TSA-Seq score percentiles (20 kb bins) for biological replicates 1 (x-axis) and 2 (y-axis) across entire percentile range (**D**) and top 10 percentile regions (**E**). Colors represent numbers of 20 kb genomic bins with given TSA-Seq score percentiles in the 2 replicates falling within given replicate 1 percentile: replicate 2 percentile histogram 2D bin (1 percentile x 1 percentile intervals). **E)** Regions between dashed lines show histogram bins in which TSA-Seq score percentiles are within a 5 percentile difference of each other in the two replicates. **F)** 4-way Venn diagram showing overlapping of SPADs across all 4 cell lines. 56.4% (129.3 Mbp) are classified as SPADs (>95th percentile) in all 4 cell lines, 12.5% (28.6 Mbp) in 3 cell lines, 13.0% in 2 cell lines (29.7 Mbp), and 18.1% (41.5 Mbp) in just 1 cell line, out of 229.1 Mbp total. However, 100% of SPADs remain near speckles (>80th percentile in relative SON TSA-Seq enrichment scores) in all 4 cell lines. **G)** Protein-coding gene FPKM in 20 SON TSA-Seq enrichment score vigintiles (division of percentile scores, 0-100, into 20 bins of 5% size each) in H1 (top left), K562 (top right), HCT116 (bottom left), and HFFc6 (bottom right). Box plots show median (inside line), 25th (box bottom) and 75th (box top) percentiles, 75th percentile to highest value within 1.5-fold of box height (top whisker), and 25th percentile to lowest value within 1.5-fold of box height (bottom whisker). **H)** Distributions of percentages (y-axis) of housekeeping (red) and non-housekeeping (blue) protein-coding genes versus SON TSA-Seq vigintiles (x-axis) in 4 cell lines.

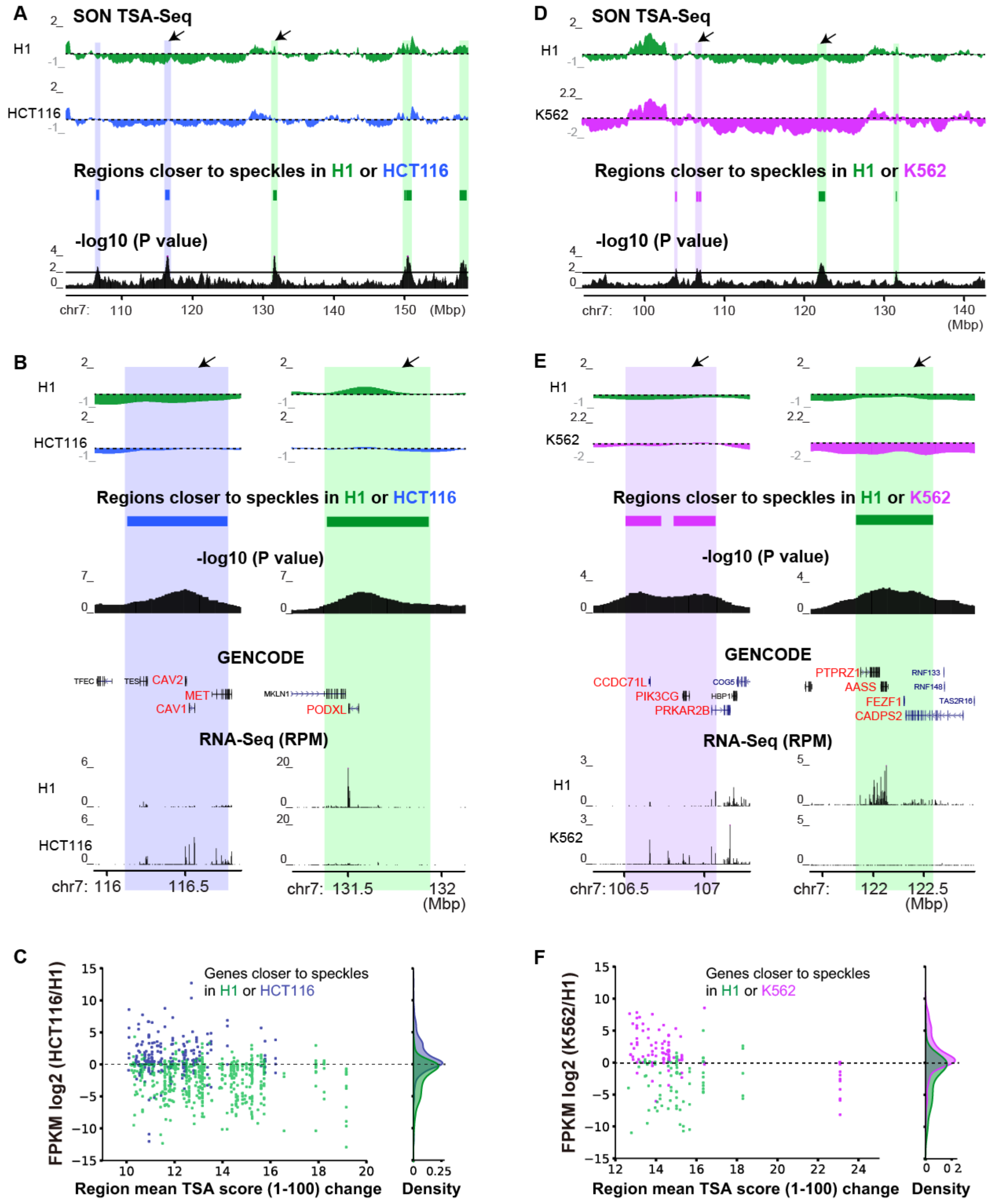


**Supplementary Figure 6. Pair-wise cell type comparisons of SON TSA-Seq mapping reveal regions with variable distances to nuclear speckles that are tightly correlated with changes in gene expression. A,**

**D)** H1 and HCT116 (**A**) or H1 and K562 (**D**) SON TSA-Seq enrichment scores (smoothed) (top, dashed lines indicate “zero” values which are regions with genome-wide average number of reads), highlighted repositioned domains (middle), and - log10 (p-value per bin for significance of change in scaled SON TSA-Seq scores) (bottom). **B, E)** Zoomed view of two regions (arrows in **A, D**) plus gene annotation (GENCODE) and RNA-Seq RPM values. **C, F)** Changes in expression of protein-coding genes in repositioned domains: scatterplots show log2 fold-changes in FPKM ratios (y-axis) between HCT116 and H1 (**C**) or between K562 and H1 (**F**) versus absolute values of changes in mean scaled TSA-Seq scores (x-axis, max-min normalized: 1-100). Gene distribution is shown as Kernel density plots on the right.

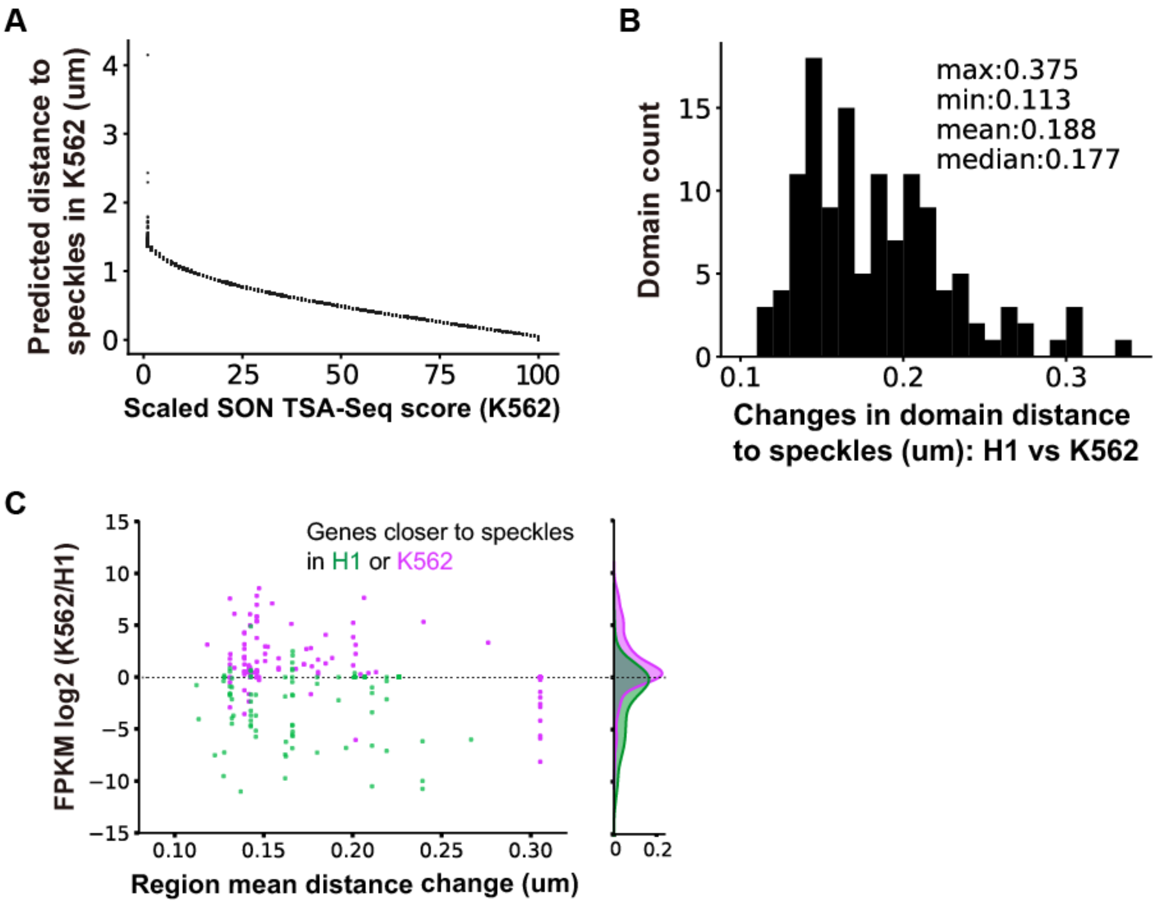


**Supplementary Figure 7. Distance calibration demonstrates small distance changes relative to speckles in H1 vs K562 comparison yet a tight correlation with changes in gene expression. A)** Scatter plot showing predicted distances in microns versus corresponding scaled TSA-Seq enrichment scores (1-100, 20kb bins). **B)** Distribution of predicted mean distance changes for each changed domain between H1 and K562 cells (x-axis: intervals of 0.01 µm). **C)** Changes in expression of protein-coding genes in repositioned domains: scatterplots show log2 fold-changes in FPKM ratios (y-axis) between K562 and H1 versus changes in mean domain distances (x-axis). Gene distribution is shown as Kernel density plots on the right.

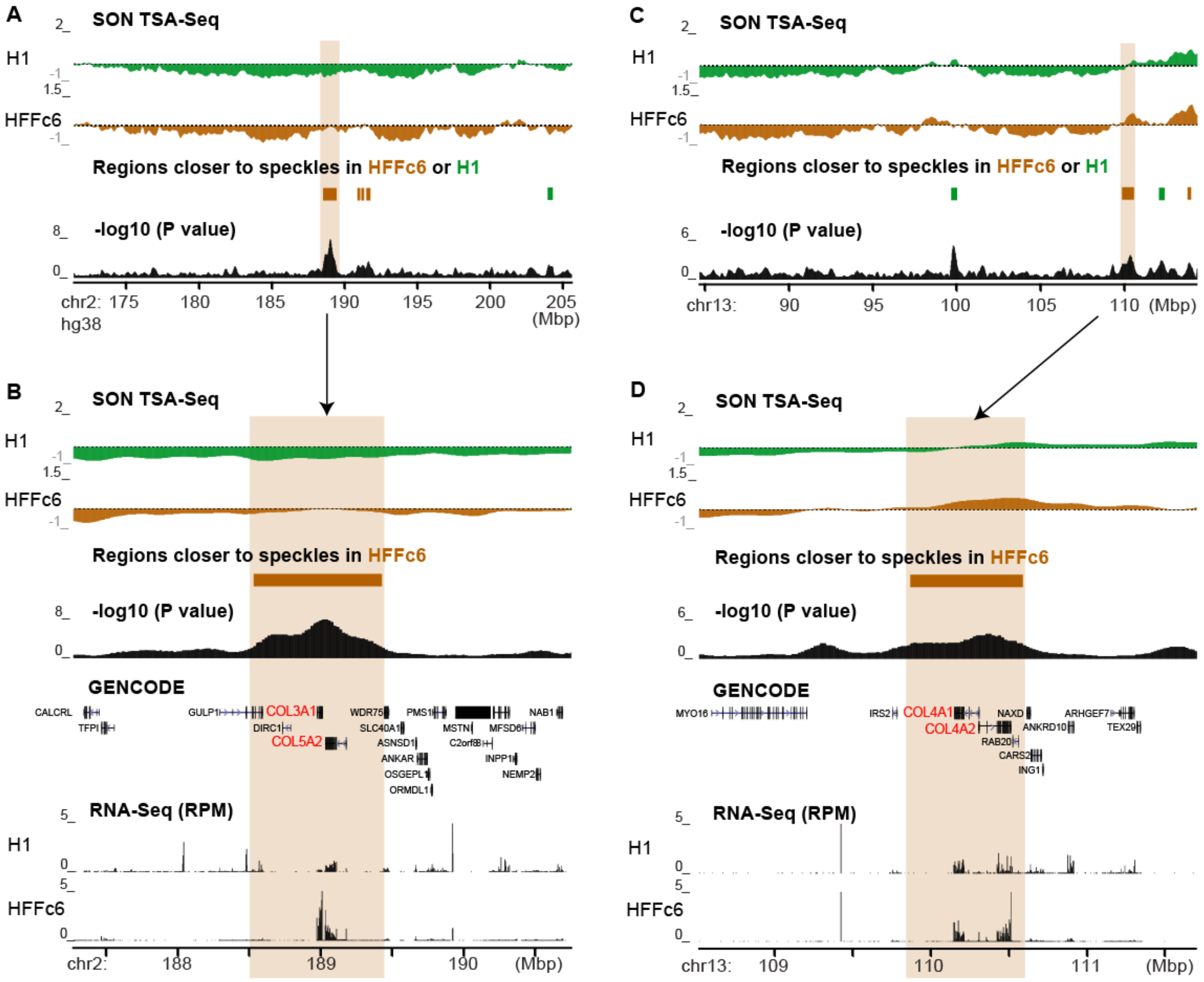


**Supplementary Figure 8. Increased Collagen gene expression in HFF fibroblasts versus H1 hESCs correlates with closer relative distance to nuclear speckles in fibroblasts versus hESCs. A, C)** Comparison of H1 and HFFc6 smoothed SON TSA-Seq enrichment scores: top track- H1 (green), middle track- HFFc6 (brown), bottom track- p-values per bin for significance of change in scaled SON TSA-Seq scores. Dashed lines (top and middle tracks) indicate "zero" values showing regions with genome-wide average number of reads. Highlighted regions (brown) below middle track containing Collagen genes (**A, B**: COL3A1, COL5A2; **C, D**: COL4A1, COL4A2) localize closer to speckles in HFFc6. **B, D)** Zoomed browser views of same highlighted regions also showing gene annotation (GENCODE v29) and RNA-Seq RPM values.

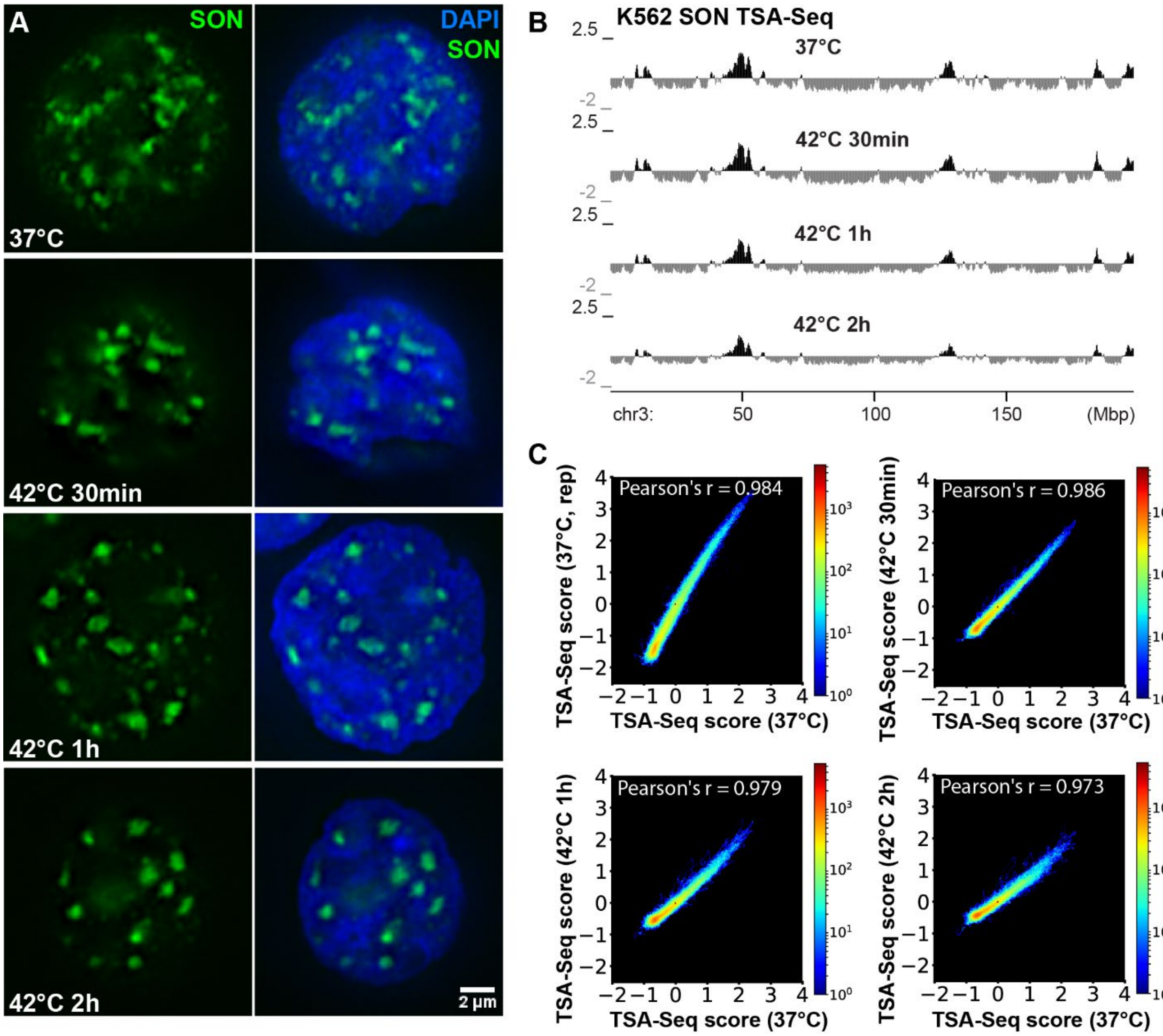


**Supplementary Figure 9. Genome organization relative to nuclear speckles as measured by TSA-Seq is largely invariant after heat shock. A)** SON immunostaining (green) in K562 cells at 37 °C or upon heat shock at 42 °C for 30 mins, 1 hr or 2hrs with DAPI (blue, right). **B)** SON TSA-Seq mapping results with or without heat shock showing smoothed TSA-Seq enrichment scores. **C)** 2D histograms showing correlation of smoothed SON TSA-Seq enrichment scores between replicates or different conditions. Colors represent numbers of 20 kb genomic bins with given TSA-Seq scores in the 2 datasets falling within given histogram 2D bin (~0.03 x 0.03 of TSA-Seq score intervals). Pearson's r: the Pearson product-moment correlation coefficient.

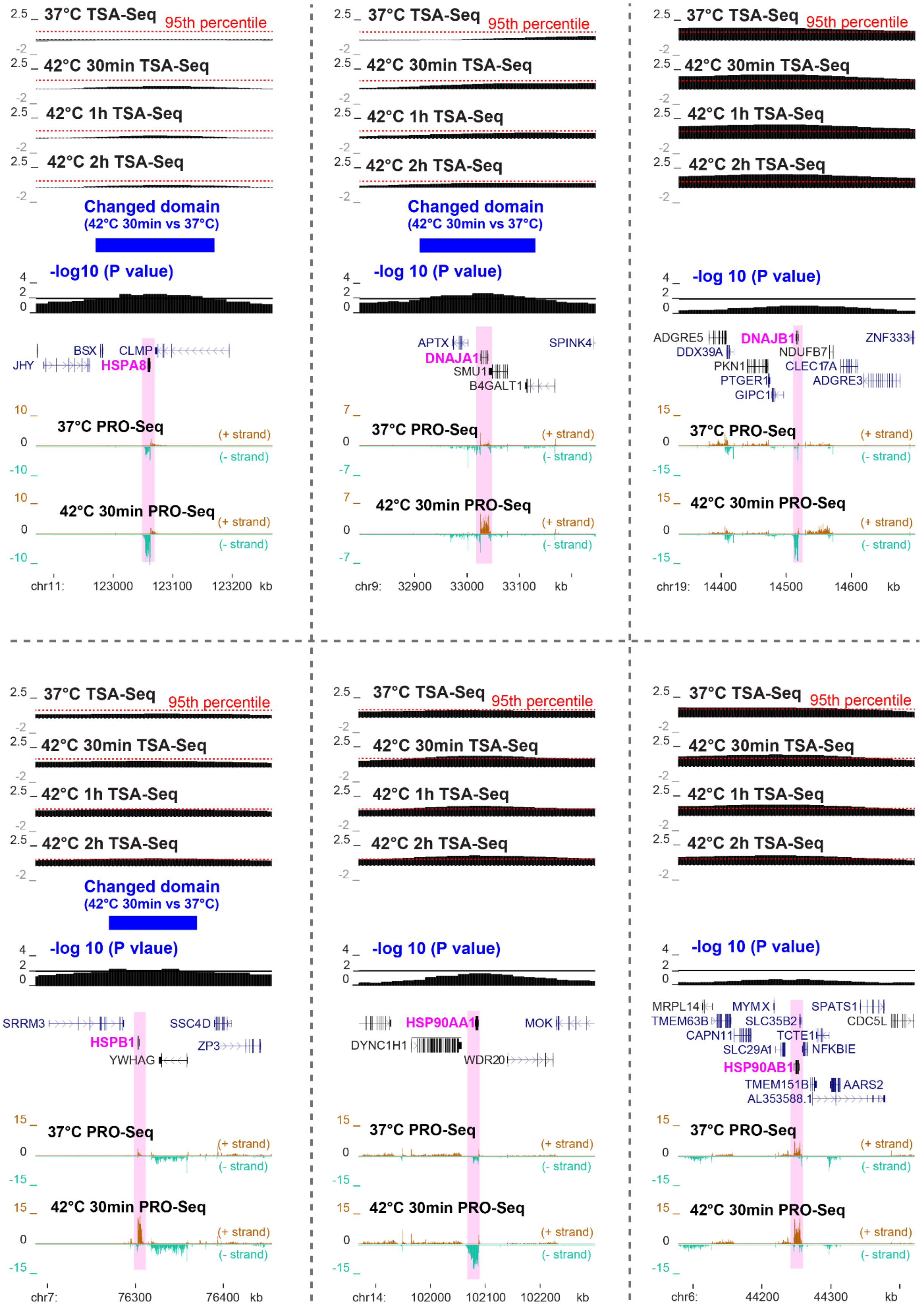

37°C TSA-Seq
95th percentile
42°C 30min TSA-Seq
42°C 1h TSA-Seq
42°C 2h TSA-Seq
Changed domain
(42°C 30min vs 37°C)
-log10 (P value)
-log 10 (P value)
-log 10 (P vlaue)
BSX
CLMP
JHY
HSPA8
37°C PRO-Seq
42°C 30min PRO-Seq
(+ strand)
(- strand)
chr11: 123000 123100 123200 kb
APTX
SPINK4
DNAJA1
SMU1
B4GALT1
chr9: 32900 33000 33100 kb
ADGRE5
DNAJB1
ZNF333
DDX39A
NDUFB7
PKN1
CLEC17A
PTGER1
ADGRE3
GIPC1
chr19: 14400 14500 14600 kb
SRRM3
SSC4D
HSPB1
ZP3
YWHAG
chr7: 76300 76400 kb
HSP90AA1
MOK
DYNC1H1
WDR20
chr14: 102000 102100 102200 kb
MRPL14
MYMX
SPATS1
TMEM63B
SLC35B2
CDC5L
CAPN11
TCTE1
SLC29A1
NFKBIE
HSP90AB1
TMEM151B
AARS2
AL353588.1
chr6: 44200 44300 kb

**Supplementary Figure 10. Major inducible heat shock protein genes reside in unchanged SPADs or domains moving small distances to speckles.** TSA-Seq and PRO-Seq profiles of multiple induced heat shock protein (HSP) genes in K562 cells upon heat shock. From top to bottom: smoothed SON TSA-Seq enrichment score tracks of control or different heat shock time points (red dashed lines indicate $95^{th}$ TSA-Seq score percentiles), significantly changed domains comparing 37 °C vs 42 °C 30 mins (blue), -log10 (p-values of 20kb-bins for statistical comparison of rescaled SON TSA-Seq scores between 37 °C and 42 °C 30 mins), gene annotation (GENCODE) and K562 PRO-Seq tracks showing normalized read count (BPM, 50 bp bins).

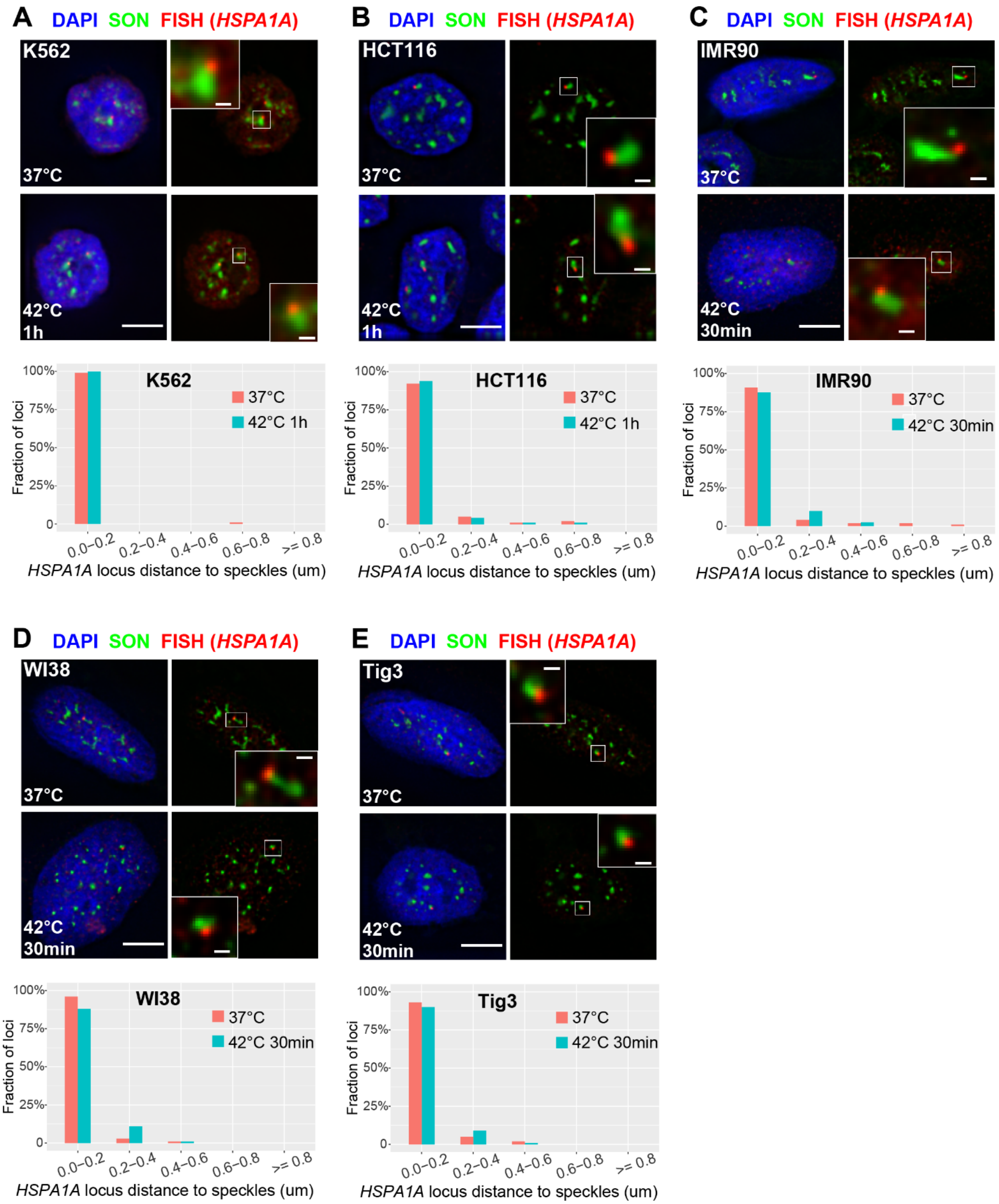


113

114 **Supplementary Figure 11.** ***HSPA1A*** **locus deterministically positions at the periphery of nuclear**

115 **speckles by microscopy in multiple cell lines. A-E) Top:** 3D immuno-FISH for *HSPA1A* locus (red) plus

SON immunostaining (green) and merged channels with DAPI (blue, left, scale bar: 5 µm) in K562 (**A**), HCT116 (**B**), IMR90 (**C**), WI38 (**D**), or Tig3 (**E**) cells at 37 °C or after 42 °C heat shock. Insets: 4× enlargement of the white-boxed image area (right, scale bar: 0.5 µm). **A-E) Bottom:** Distribution comparison of distances from *HSPA1A* FISH signal to a closest nuclear speckle in K562 (**A**), HCT116 (**B**), IMR90 (**C**), WI38 (**D**), or Tig3 (**E**) cells between control and heat shock treated samples. K562: n=101 (37 °C) or 104 (42 °C 1hr). HCT116: n=101 (37 °C) or 97 (42 °C 1hr). IMR90: n=98 (37 °C or 42 °C 30 mins). WI38: n=100 (37 °C or 42 °C 30 mins). Tig3: n=100 (37 °C or 42 °C 30 mins).

### Supplementary tables

#### Supplementary Table 1

| BAC | Genome coordinates (hg38) | Mean distance to speckles (um) |
|---|---|---|
| RP11-634L10 | chr17:81,838,939-82,011,417 | 0.09 |
| RP11-479I13 | chr6:31,726,514-31,941,167 | 0.11 |
| RP11-264N5 | chr7:100,470,712-100,665,336 | 0.16 |
| RP11-1058N17 | chr18:48,801,893-48,998,009 | 0.47 |
| CTD-3106L12 | chr2:24,775,317-24,986,884 | 0.5 |
| RP11-997B19 | chr17:71,701,964-71,881,248 | 0.81 |
| RP11-978O5 | chr2:22,703,020-22,897,142 | 0.97 |
| RP11-846O11 | chr18:41,032,947-41,237,108 | 0.98 |
| RP11-302K17 | chr10:102,058,244-102,216,677 | 0.25 |
| RP11-246J15 | chr1:202,111,935-202,271,377 | 0.36 |
| CTD-3244P16 | chr10:102,990,564-103,165,048 | 0.47 |
| CTD-2503D10 | chr10:103,950,393-104,169,256 | 0.51 |
| RP11-729K13 | chr2:30,387,390-30,582,345 | 0.6 |
| RP11-53I19 | chr6:23,302,021-23,446,224 | 0.76 |
| RP11-543G21 | chr1:199,421,727-199,594,861 | 0.92 |
| RP11-1047B3 | chr7:114,496,093-114,692,887 | 0.97 |

#### Supplementary Table 2

| K562 SON TSA | Condition A | Condition B | Condition C | Condition D | Condition E |
|---|---|---|---|---|---|
| $\boldsymbol{y_0}$ | 0.27 | 0.35 | 0.24 | 0.38 | 0.28 |
| $\boldsymbol{A}$ | 9.69 | 6.10 | 7.7 | 5.08 | 6.75 |
| $\boldsymbol{R_0}$ | -4.28 | -3.53 | -3.80 | -3.43 | -3.79 |

**Supplementary Table 3**

| Cell line | Organism | Lineage |
|---|---|---|
| **H1** | Human | Embryonic stem cell |
| **K562** | Human | Erythroleukemia |
| **HCT116** | Human | Colorectal carcinoma (epithelial) |
| **HFFc6** | Human | Foreskin fibroblast |

**Supplementary Table 4**

| ***HSPH1* smRNA-FISH probes** |
|---|
| agaatctgcggcatttactc |
| tggctgataagaaaccctgg |
| tccaatttgtgatggcaagc |
| cgtgaatccctgagacaatc |
| cccagtaggttttgaaaacc |
| gcgcaaagatatgcgcgaag |
| cggaagaaaaaggccggcag |
| cttccagttcatttcgagat |
| tggacaagcagcgacacaag |
| tggtcgctaacaacaatccg |
| gtgggctaatctttcatagg |
| ggccaacaatgttagcaacg |
| catctttcgacaatgatccc |
| gcgtcctagcacaaaagtat |
| ttagttcagtgacgacatcc |
| taattgtattaccccagtca |
| tcataaacgttctcagcacc |
| acagtcttcctaacgttact |
| aacttagatctgcaggggta |

| |
|---|
| aaggggaactccaatactgc |
| atctactcattttcctaggg |
| gcatcactgtaagctctgaa |
| tacctgcttaggtatgagta |
| ccgcacaataacagagcaga |
| gtcgagggaaaggagtgact |
| tttgttctgatataccaccg |
| gatgtgactctacctttcag |
| ttgaccctacacacgactaa |
| tactaccccaaaaacctctg |
| aggggatagggaacaagcat |

**Supplementary Table 5 is a separate Excel file showing public RNA-Seq datasets used for correlation with TSA-Seq and for differential expression analysis between H1 and HFF cells.**

**Supplementary Table 6 is a separate Excel file showing results of RNA-Seq data analysis and correlation TSA-Seq data in the four cell lines.**

**Supplementary Table 7 is a separate Excel file showing all identified differentially expressed protein-coding genes between H1 and HFF cells.**

**Supplementary Table 8 is a separate Excel file showing results of gene ontology analysis for differentially expressed protein-coding genes in regions that are relatively closer to speckles versus that are not relatively closer to speckles between H1 and HFF cells.**

**Supplementary Table 9 is a separate Excel file showing summary of TSA-Seq data used in this study.**
